# Rapid expansion and visual specialization of learning and memory centers in Heliconiini butterflies

**DOI:** 10.1101/2022.09.23.509163

**Authors:** Antoine Couto, Fletcher J Young, Daniele Atzeni, Simon Marty, Lina Melo-Flórez, Laura Hebberecht, Monica Monllor, Chris Neal, Francesco Cicconardi, W Owen McMillan, Stephen H. Montgomery

**Author notes:** contributed equally (Antoine Couto, Fletcher J Young).

## Abstract

How do neural systems evolve to support new behaviors? Changes in the abundance and diversity of neural cell types, and their connectivity, shape brain composition and provide the substrate for behavioral variation. We describe a striking example of neural elaboration in an ecologically diverse tribe of Heliconiini butterflies. By building extensive new datasets of neural traits across the tribe, we identify major bursts in the size and cellular composition of the mushroom bodies, central brain structures essential for learning and memory. These expansion events are associated with increased innervation form visual centers and coincide with enhanced performance in multiple cognitive assays. This suite of neural and cognitive changes is likely tied to the emergence of derived foraging behaviors, facilitated by localized specialization of neural networks.

**One-Sentence Summary:** Major shifts in brain composition and behavior in butterflies with unique foraging and dietary behaviors.

## Introduction

Evolutionary innovation, elaboration and refinement of neural systems underpins the diversity of behavior, sensory and cognitive abilities that we see across animal life. Models of brain evolution increasingly emphasize the coordinated evolution of functionally related networks, but with localized specialization where selection targets refinement of existing processes (*1–3*). Under this model, increased behavioral precision, or diversification, can occur through replication of established cell types and circuits within generally conserved networks (*4, 5*). While existing data provide patterns of variation consistent with this process, demonstrating associations between specialized refinements of specific neuronal circuits and behavioral enhancement remains challenging.

Cases of ecological innovation can provide unique opportunities to link neural and behavioral evolution (*6, 7*) particularly when grounded in robust phylogenetic frameworks. For example, uniquely among butterflies, adult *Heliconius* actively collect and digest pollen (*8, 9*), providing an adult source of essential amino acids (*8, 10*) and facilitating a greatly extended reproductive lifespan (*11*). This dietary innovation is accompanied by the evolution of trap-line foraging, where individuals learn foraging routes between resources with high spatial and temporal fidelity (*12–14*). Among insects, this foraging strategy is also found among species of bee (*15, 16*), and is thought to rely on visual memories and landmark cues (*17*). This suggests that the evolution of pollen feeding in *Heliconius* may have required neural and cognitive enhancements to support optimal foraging for low-density pollen resources through enhanced spatial memory. Indeed, preliminary evidence suggests that *Heliconius* have expanded mushroom bodies, an insect learning and memory center (*18*), relative to their non-pollen-feeding relatives in the Heliconiini tribe (*19, 20*). While not essential for spatial memory in *Drosophila* (*21*), mounting evidence from empirical (*22–24*) and theoretical modelling (*25, 26*) implicates these structures in spatial orientation in other insects. However, knowledge of how the adaptative evolution of sensory integration and learning centres supports cognitive enhancement is limited. Here, we provide an extensive analysis of mushroom body expansion in *Heliconius*, with new, phylogenetically dense data across all major Heliconiini lineages. By incorporating multiple neuroanatomical and behavioral traits, we establish a rich system to link neural elaboration and behavioral innovation.

## i. Massive, independent evolution in Heliconiini mushroom body size

We generated an extensive dataset of brain composition for 318 wild-caught individuals from 41 Heliconiini species (average 8 individuals/species), including 30 species and sub-species of *Heliconius* and representatives from each Heliconiini genus (Figure 1). 3D volumetric reconstructions of immuno-stained whole brains revealed dramatic variation in mushroom body size across the tribe. In raw volume, mushroom bodies vary by 26.5-fold, from ~3×10^6^ μm^3^ to ~70×10^6^ μm^3^ per hemisphere (Table S1). This level of variation is unmatched by major visual or olfactory brain structures (medulla: 7.9-fold variation; antennal lobe: 6.2-fold variation), or the remaining volume of the central brain (7.8-fold variation). Across all individuals, correcting for allometric effects and phylogenetic relatedness using MCMCglmm (*27*), mushroom body volume is associated with variation in both visual and olfactory structures (medulla: p_MCMC_ = 0.002; antennal lobe: p_MCMC_ < 0.001). However, significant variation between phylogenetic groups still persists, with *Heliconius* having significantly larger mushroom bodies compared to other genera (p_MCMC_ < 0.001). This is not the case for major visual or olfactory brain regions (medulla: p_MCMC_ = 0.482; antennal lobe: p_MCMC_ = 0.202; Figure 2E-H). Repeating this analysis including more narrow phylogenetic groupings (outgroup genus, or *Heliconius* subclades) suggests that variation in mushroom body size is not distributed bimodally, but varies both within *Heliconius* and across the outgroup genera (Supplementary Results, Figure S3, Tables S8–9). Interestingly, we also identify a *Heliconius* specific effect of sex (interaction *Heliconius*\*Sex p_MCMC_ < 0.001), with females having larger mushroom bodies on average than males (p_MCMC_ < 0.001; Figure S2), which could reflect sex differences in foraging for pollen or host plant resources (*28*).

**Figure 1.**
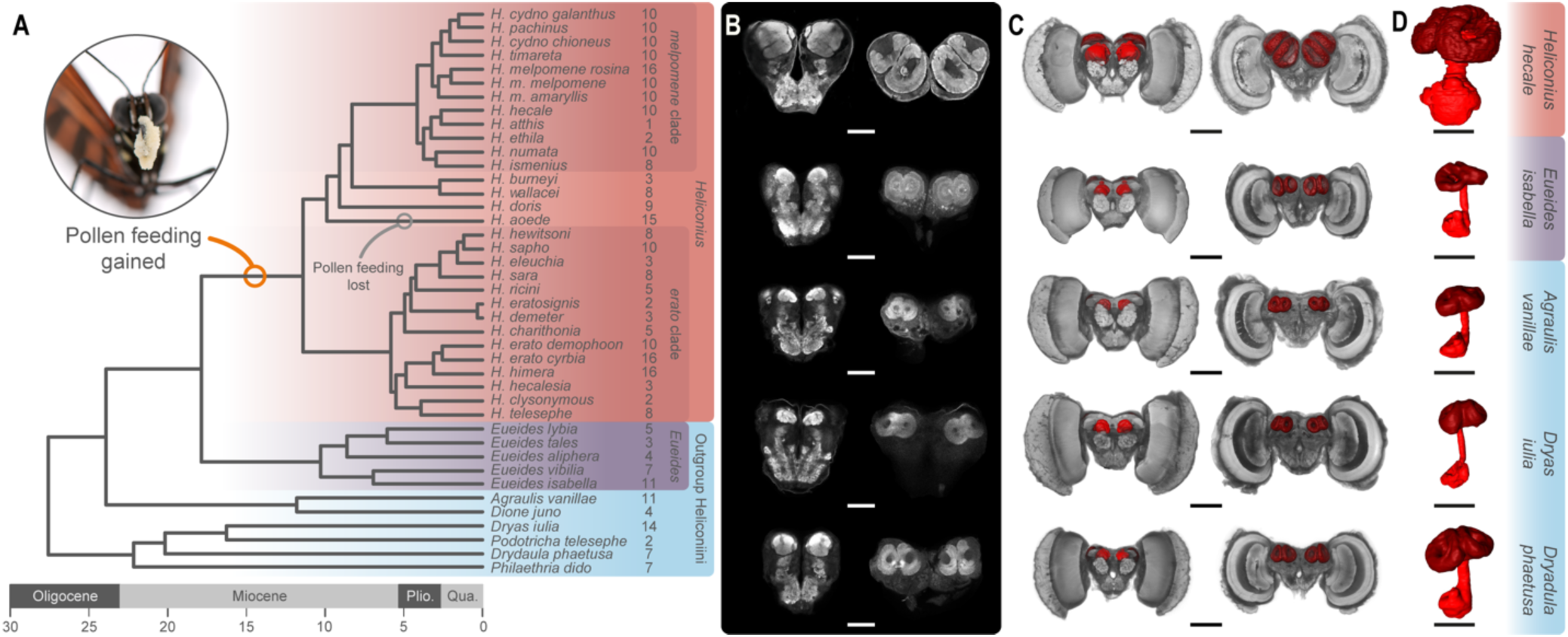
Variation in absolute mushroom body size across the Heliconiini. (**A**) Dated phylogeny showing the Heliconiini species sampled and the major clades. **(B, C)** Selected neuroanatomical detail from five species: (**B**) Confocal scans showing the anterior (left) and posterior (right) of the central brain (scale bars = 250 μm). (**C**) 3D reconstructions of the whole brain with anterior (left) and posterior (right) views showing the mushroom body (red) (scale bars = 500 μm). (**D**) Isolated 3D reconstructions of the mushroom body, showing the calyx (dark red), peduncles and lobes (light red) (scale bars = 250 μm).

**Figure 2.**
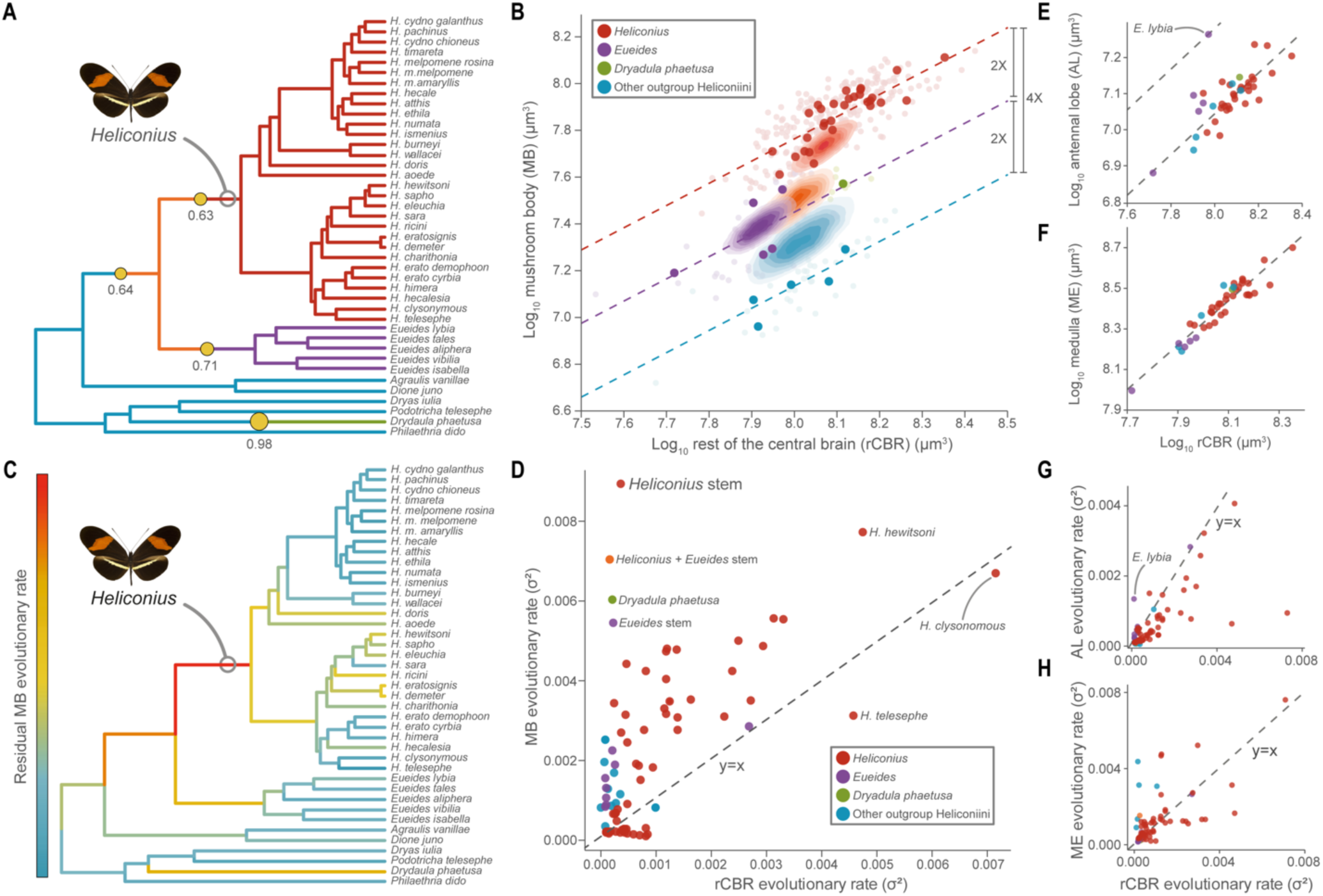
Major shifts in the relative size and evolutionary rate of the mushroom body in *Heliconiini* butterflies. (**A** and **B**) Phylogenetic shifts in the scaling relationship between the volume of the mushroom body (MB) and the rest of the central brain (rCBR) across 41 Heliconiini taxa (posterior probability > 0.5). Relative to outgroup Heliconiini (blue), MB volumes are twice as large in *Eueides* (purple) and *Dryadula phaetusa* (green) and four times as large in *Heliconius* (red). Solid points = species means; faded points = individuals. Estimated ancestral states for each group shown by density maps. (**C** and **D**) The branch leading to *Heliconius* shows a marked increase in the evolutionary rate of MB volume, with a slightly less elevated rate along the branch leading to *Heliconius*+*Eueides*. (**E**-**H**) Shifts in MB size and evolutionary rate are not reflected in either the antennal lobe or the medulla.

## ii. Multiple bursts of accelerated rates of mushroom body expansion

To further explore the evolutionary history of mushroom body size across the Heliconiini, we used multiple methods to reconstruct ancestral states, evolutionary rates and shifts in allometric scaling. First, we used BAYOU (*29*) to identify shifts in scaling between mushroom body size and central brain size. The best fitting model permitted shifts in elevation specifically (marginal likelihoods: elevation shifts = 54.252; slope and elevation shifts = 37.093; none = 32.556) and identified four shifts with a posterior probability greater than 0.5 (Figure 2A), representing increases in relative mushroom body size at the internal branches leading to *Heliconius+Eueides* (post. prob. = 0.64) and *Heliconius* (post. prob. = 0.63), as well an independent expansion in *Dryadula* (post. prob. = 0.98), and a reduction at the base of *Eueides* (post. prob. 0.71). These shifts result in phylogenetic groupings of species with approximately common allometric scaling of the mushroom body, with convergent allometries between *Eueides* and *Dryadula*, which are intermediate between *Heliconius* and all other outgroup genera (Figure 2B). These results are supported by pairwise comparisons among all genera (Figure S3, Tables S8–9), and by ancestral state reconstructions which again imply serial shifts in mushroom body size independently of central brain size, culminating in *Heliconius,* where the estimated ancestral state is within the range of extant *Heliconius* species (Figure 2B).

We next used evolutionary models that permit branch-specific shifts in rate parameters to estimate points in the Heliconiini phylogeny where either mushroom body or central brain size evolved rapidly. Two alternative methods (*30, 31*) (Figure 2C, D; Figure S5) identify high rates of evolution for mushroom body size specifically, on multiple internal branches (e.g. Figure 2C, D). The highlighted branches are the same as in the BAYOU analysis, with the stem *Heliconius* branch standing out as having the highest rate of evolution in mushroom body size relative to central brain size (Figure 2D). Analyzing sensory brain regions in the same way fails to recapitulate these shifts and bursts in evolutionary rate (Figure 2E-H), demonstrating that increases in mushroom body size are not primarily caused by changes in the sensory periphery. Finally, to contextualize this variation within a broader sample of butterflies, we used the same approach to reanalyze a phylogenetically broad dataset of 41 species of North American butterflies which includes *Heliconius charithonia* and the non-pollen feeding Heliconiini *Agraulis vanillae* (*32*). Both the BAYOU and rates analysis highlight the *Heliconius* branch as the sole stand-out lineage, and a remarkably clear outlier in mushroom body evolution across butterflies (Figure S4).

## iii. Volumetric expansion is closely tied to increases in Kenyon cells

These volumetric expansions could be due to a combination of i) increases in the number of Kenyon cells, the intrinsic mushroom body neurons; and/or ii) increases in the synaptic contacts (branching patterns) made by Kenyon cells, which may result in altered volumes of calyx per Kenyon cell, or increased synapse density or number. To address this, we estimated total Kenyon cell number by staining samples with neural and nuclear markers to measure the volume of the Kenyon cell cluster, which surrounds the posterior calyx (Figure 3A), and the density of nuclei within (Figure 3A-C). We did this for three *Heliconius* and three outgroup Heliconiini species. Our estimates of Kenyon cell number per hemispheres vary from ~11,000 in *Agraulis vanillae* to ~80,000 in *Heliconius hortense* (Table S2). Total Kenyon cell number varies significantly across species (χ^2^ = 1475.3, d.f. = 5, p < 0.0001; Figure 3D; Table S2), and is significantly higher in *Heliconius* (χ^2^ = 44.83, d.f. = 1, p < 0.0001). Kenyon cell density varies across species (χ^2^ = 34.775, d.f. = 5, p < 0.0001), however it is does not differ consistently between *Heliconius* and other Heliconiini overall (χ^2^ = 0.3617, d.f. = 1, p = 0.548). We also independently verified these estimates by counting cross sectioned Kenyon cell axons running through the mushroom body peduncle in three species (n=2/species), imaged using electron microscopy (Figure S6). This produced estimates of ~13,000 Kenyon cells in *Dryas iulia*, ~52,000 in *Heliconius erato*, and ~78,000 in *Heliconius melpomene*, which are within the range of estimates produced using immunostaining.

**Figure 3.**
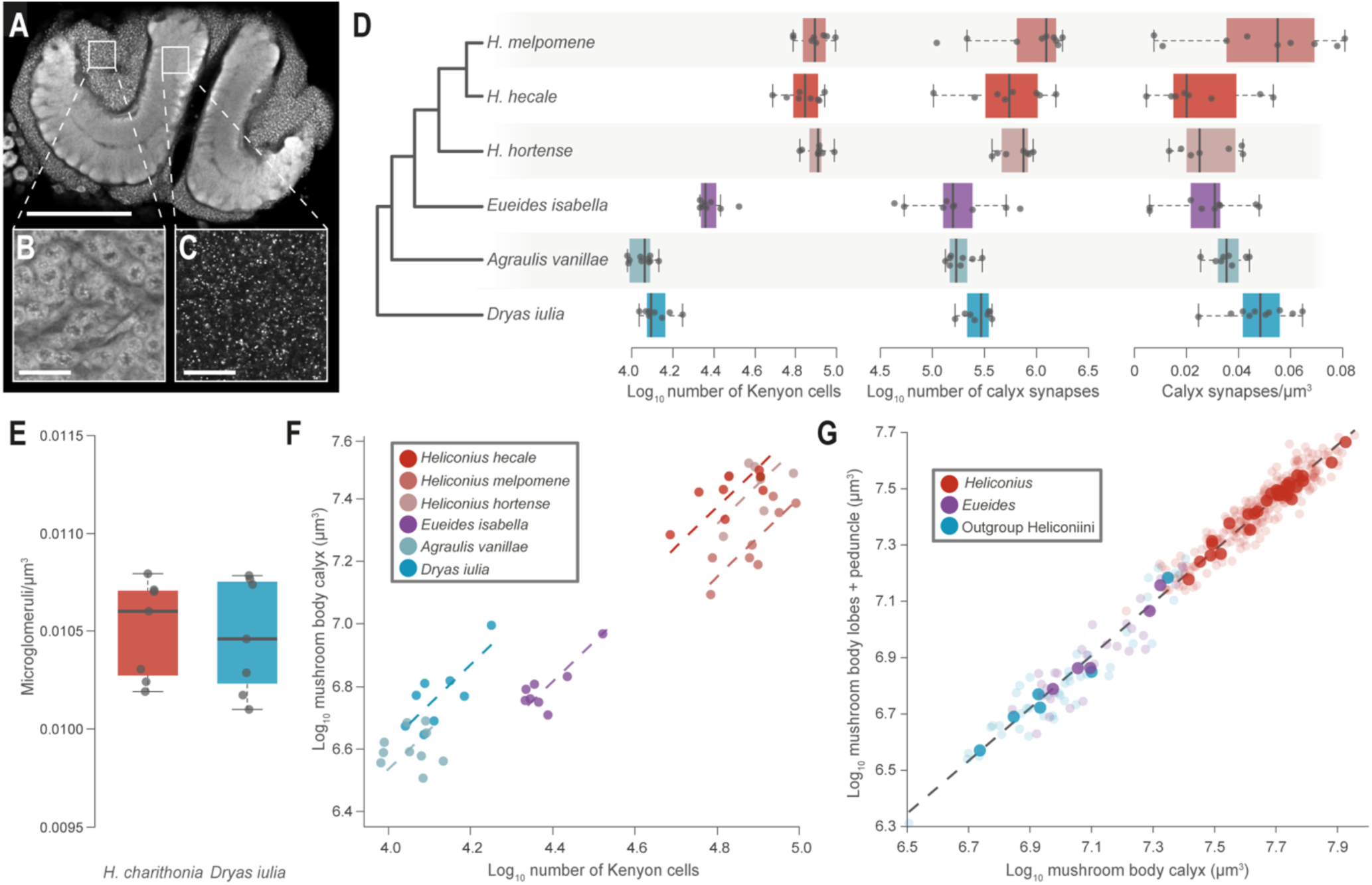
Increased Kenyon cells and synapses numbers in *Heliconius*, but conserved cellular scaling. (**A**) *Heliconius* mushroom body (scale bar = 200 μm), showing (**B**) the intrinsic neurons, the Kenyon cells (KCs) (scale bar = 50 μm) and (**C**) synapses (scale bar = 50 μm). (**D**) *Heliconius* show increases in the number of Kenyon cells and synapses, but not synapse density. (**E**) Microglomeruli density does not differ between *H. charithonia* and *Dryas iulia*. (**F**) The relationship between calyx volume and KC number varies between species, but not overall between *Heliconius* and other Heliconiini. (**G**) The relationship between the major components of the mushroom body (the calyx and the lobes and peduncle) is conserved across the Heliconiini. Solid points = species means; faded points = individuals.

Scaling between the calyx volume and total Kenyon cell number is significantly different across species (χ^2^ = 90.749, d.f. = 5, p < 0.0001), resulting in elevation shifts (*W* = 90.06, d.f. = 5, p < 0.0001). Post-hoc pairwise analysis indicates that this is primarily due to a shift between *Heliconius+Eueides* and other genera, which results in a smaller volume of calyx per Kenyon cell (Figure 3F). We also quantified the density of synapses within the calyx to indirectly test for evidence of altered connectivity. Although we do find an effect of species on synapse density (χ^2^ = 17.846, d.f. = 5, p = 0.003), there is no evidence that *Heliconius* consistently differ from other genera (χ^2^ = 0.077, d.f. = 1, p = 0.782) although, due to the increased cell number and calyx volume, they do have significantly higher total synapse estimates (χ^2^ = 32.664, d.f. = 1, p < 0.0001). We supplemented these data with estimates of microglomeruli (a synaptic complex (*33*); Figure S7) density in *Heliconius charithonia* and *Dryas iulia*, which also provides no evidence for a shift in density (χ^2^ = 0.055, d.f. = 1, p = 0.815; Figure 3E). Finally, we explored whether the scaling relationship between the calyx, where Kenyon cells synapse with incoming projection neurons, and the lobes, formed by the main axon terminals of Kenyon cells and the dendrites of mushroom body output neurons, is altered in species with expanded mushroom bodies. We found a remarkably consistent scaling relationship across all Heliconiini, with no shift in *Heliconius* (p_MCMC_ = 0.180; Figure 3G).

Together, we propose these results are consistent with mushroom body expansion being produced by increased Kenyon cell production and a replication, rather than innovation, of neural circuitry. Theoretical modelling of mushroom body circuitry suggests that, under conserved levels of average Kenyon cell activity, the capacity to store visual patterns increases logarithmically with Kenyon cell number (*25*). Our data therefore indicate that *Heliconius* likely have the capacity to store many more engrams, a unit of cognitive information imprinted in neural systems (*34*), than the ancestral Heliconiini, providing a substrate for improved navigational performance through memorization of landmarks as required for trap-line foraging.

## iv. Mushroom body expansion is associated with increased visual input

While our data currently imply replication of conserved internal circuitry, the possible link between mushroom body expansion and visually-orientated spatial memory would predict a degree of visual specialization of mushroom body function. To test this, we first confirmed that visual brain regions send projections to the mushroom body by injecting fluorescent retrograde tracers into the calyx. This revealed two major origins of projection neurons, the antennal lobe and the optic lobe (Figure S8). In the optic lobe, staining was diffuse across multiple structures, but concentrated in the ventral lobula, which we suggest may act as a relay center to the central brain (Figure S8). Notably, this structure is also highly variable in size (31.6-fold variation), but is not specifically expanded in *Heliconius* (p_MCMC_ = 0.482; Figure S9). We next differentially traced sensory projections from visual and olfactory neuropils to record the location and volume of calyx receiving input from each sensory modality in eight species, including four *Heliconius* and four outgroup Heliconiini. In all species, the calyx is topographically segregated by sensory input (Figure 4C-H), as has been observed in some other butterflies (*35*), enabling segmentation of discrete areas of calyx receiving visual or olfactory input. The volume of both the olfactory (χ^2^ = 10.396, d.f. = 1, p = 0.0012. Figure 4I) and visual (χ^2^ = 33.8, d.f. = 1, p < 0.0001, Figure 4I) calyx are expanded in *Heliconius*. However, variation in the visual calyx volume (Cohen’s d = −10.1) is considerably larger than for the olfactory calyx (Cohen’s d = −3.24, Figure 4I). The scaling relationship between these two regions is also significantly different across species (χ^2^ = 700.74, d.f. = 7, p < 0.0001), with BAYOU identifying a major shift between *Heliconius* and other genera (post prob = 0.7; Figure 4I). This results in *Heliconius* having visual calyx volumes ~2-fold larger than would be predicted by olfactory calyx volume. Among the outgroup genera we also observe post-hoc shifts in scaling that may suggest increased visual input also contributes to smaller scale variation in mushroom body expansion (Table S13–14). However, the shift in *Heliconius* is more extreme than this pattern would predict (Figure 4J) and demonstrates that a major change in the degree of visual processing in the mushroom bodies coincided with the origin of pollen feeding.

**Figure 4.**
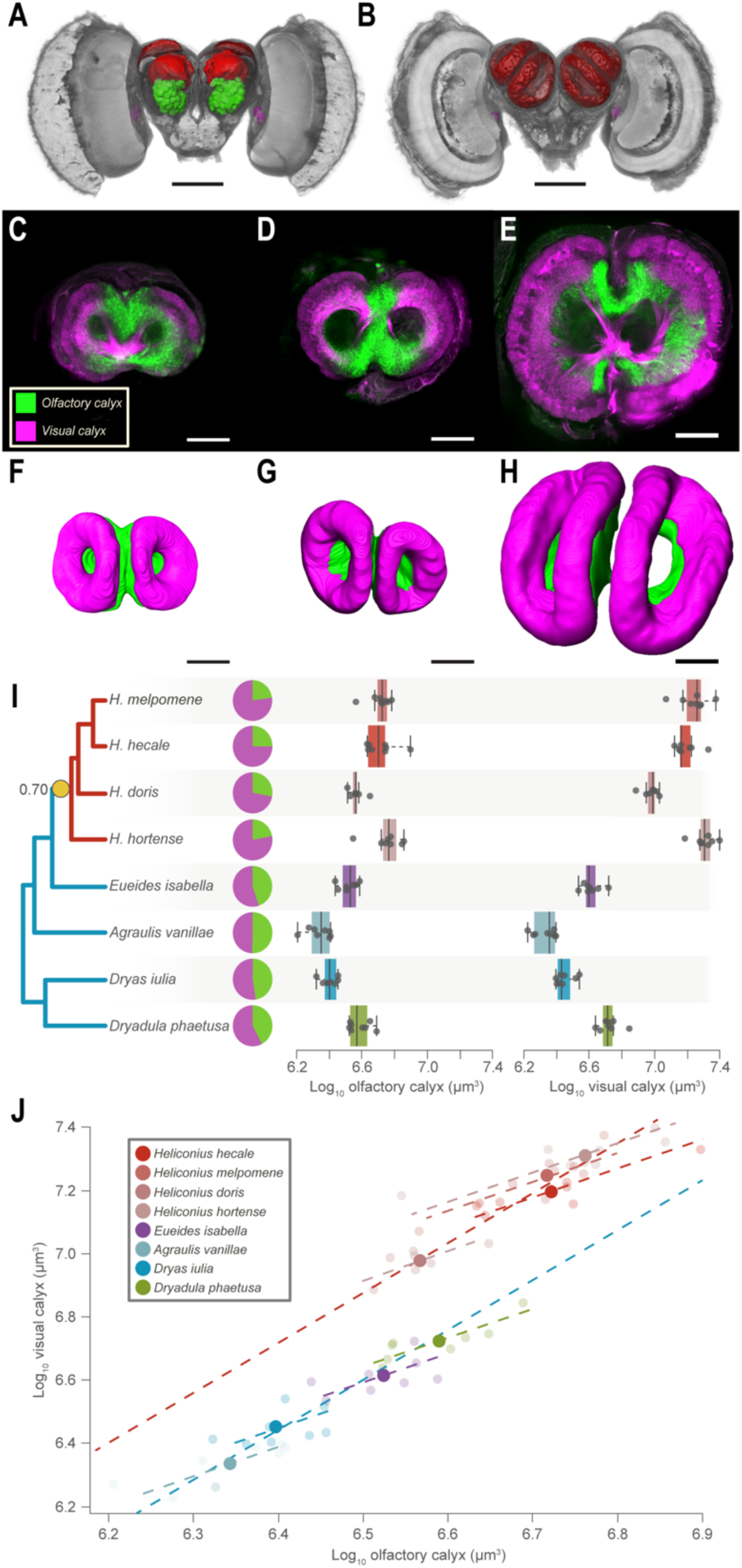
Mushroom body expansion in *Heliconius* driven primarily by increased visual input. (**A**) Anterior and (**B**) posterior 3D-reconstructions of *H. hecale* brain showing injection sites for the tracing of sensory projections to the MB (red neuropil). Olfactory projections neurons were traced from the antennal lobe (green neuropil) while visual projection were traced from injections around the ventral lobula (magenta) (scale bars = 500 μm). (**C**) Optical section of the mushroom bodies acquired with a confocal microscope in *Dryas iulia*, (**D**) *Eueides isabella* and (**E**) *H. hecale.* Inputs from olfactory (green) and visual (magenta) sensory neuropils terminate in segregated regions of the calyx. (**F**-**H)** 3D volumetric reconstructions (scale bars = 100 μm) of the visual and olfactory regions of the calyx in (**F**) *Dryas iulia*, (**G**) *Eueides isabella* and (**H**) *H. hecale*. (**I**) *Heliconius* exhibit a volumetric increase in the olfactory region of the mushroom body, but a greater increase in the visual region. (**J**) Controlling for olfactory calyx volume, *Heliconius* exhibit an upshift in size of the visual calyx. Solid points = species means; faded points = individuals.

## v. No single ecological explanation for mushroom body expansion

Increased visual processing is consistent with a plausible link between mushroom body expansion and visually-orientated spatial memory. However, our comparative analysis reveals multiple bursts in mushroom body size across the phylogeny. Ancestral state reconstructions of the presence/absence of pollen feeding imply a single origin for this trait at the base of *Heliconius*, with a secondary loss in *H. aoede* (Figure S10). The gain of pollen feeding coincides with the highest rate of mushroom body evolution (Figure 2, Figure S5) and the dominant shift in visual innervation to the calyx (Figure 4J) strongly implicating this innovation as a causative agent in mushroom body expansion. However, the presence of pollen feeding alone (DIC = −840.512) does not fit the data better than a general genus effect (DIC = −840.892), and indeed we do not identify a secondary reduction in mushroom body size in *H. aoede*, the sole *Heliconius* lineage to have lost pollen feeding (Figure S3; pMCMC = 0.543). This suggests pollen feeding does not explain all variation in mushroom body size across the tribe. We therefore explored whether additional traits that could plausibly be linked to mushroom body function can also be linked to variation in size across the tribe. First, Heliconiini share larval hostplants from the Passifloraceae, and levels of hostplant generalism could explain some variation in mushroom body size and plasticity (*36, 37*). However, using a dataset of hostplant records (*38*) we found no association between the number of hostplants used and mushroom body size (pMCMC=0.492). Second, some Heliconiini form aggregated roosts at night, and it has been argued that this could facilitate social transfer of information on resource location (*39*) and exert distinct selection pressures on the processing of conspecific cues in social species. However, we again found that the degree of social roosting had no power to explain variation in mushroom body size (pMCMC=0.857). One outstanding explanation is that the cognitive processes required for trap-line foraging are co-opted or refined from those supporting other, related, behaviors. For example, *Heliconius* have strong site fidelity and homing ability (*40*), a likely pre-requisite for trap-lining, but also a trait shared with at least some territorial species of *Eueides* (*41*). True site fidelity likely requires a degree of spatial memory, and we suggest variation in this function may explain independent shifts in mushroom body size, providing the foundation for the extreme expansion observed in *Heliconius*. Unfortunately, current ecological data on the movement ecology of non-*Helconius* Heliconiini is limited, prohibiting formal tests of this hypothesis.

## vi. Evidence for shifts in multiple cognitive assays in *Heliconius*

Nevertheless, there is a clear prediction that increases in mushroom body size should be associated with enhanced performance in some visual learning and memory contexts (*25*). *Heliconius* are capable of associative learning (A+, B−) between a reward and either color (*42–44*), shape (*45*), or odour cues (*44*), as well as contextual cues such as time of day (*43*). Natural foraging, however, likely involves encounters with complex combinations of cues. We therefore focused on two “non-elemental” learning tasks (*46*); positive pattern learning (A-, B-, AB+) and biconditional discrimination (AB+, CD+, AC-, BD-). Using artificial, colored feeders, we trained individuals of *Dryas iulia* and *Heliconius erato*, as representatives of species with small and large mushroom bodies respectively, to solve these tasks in insectary conditions (Supplementary Methods). We found that both *D. iulia* (Z ratio = −9.182, p < 0.0001) and *H. erato* (Z ratio = −16.396, p < 0.0001) can solve positive patterning tasks. However, a significant interaction between species and trial indicates that *H. erato* are more accurate in their post-training performance (χ^2^ = 66.533, d.f. = 1, p < 0.0001; Figure 5A). In the more challenging biconditional discrimination task this interaction is also found (χ^2^ = 20.727, d.f. = 1, p < 0.0001) with *H. erato* (Z ratio = −5.465, p < 0.001, but not *D. iulia* (Z ratio = −1.241, p = 0.601), being able to learn these complex cue combinations (Figure 5B). Configural learning, particularly the integration of multisensory cues, plays a crucial role in insect navigation (*24, 47*), and these results suggest that mushroom body expansion in *Heliconius* may support improvements in this ability. Computational models of mushroom body circuitry suggest solutions for these more complex tasks can emerge from circuits supporting simple associative learning (A+, B-), and that synaptic plasticity between projection neurons and Kenyon cells may be critical in mediating this process (*48*). If so, our results may suggest a potential shift in synaptic plasticity accompanied mushroom body expansion, or that the increases in the amount of visual projection or calycal synapses affects this computation.

**Figure 5.**
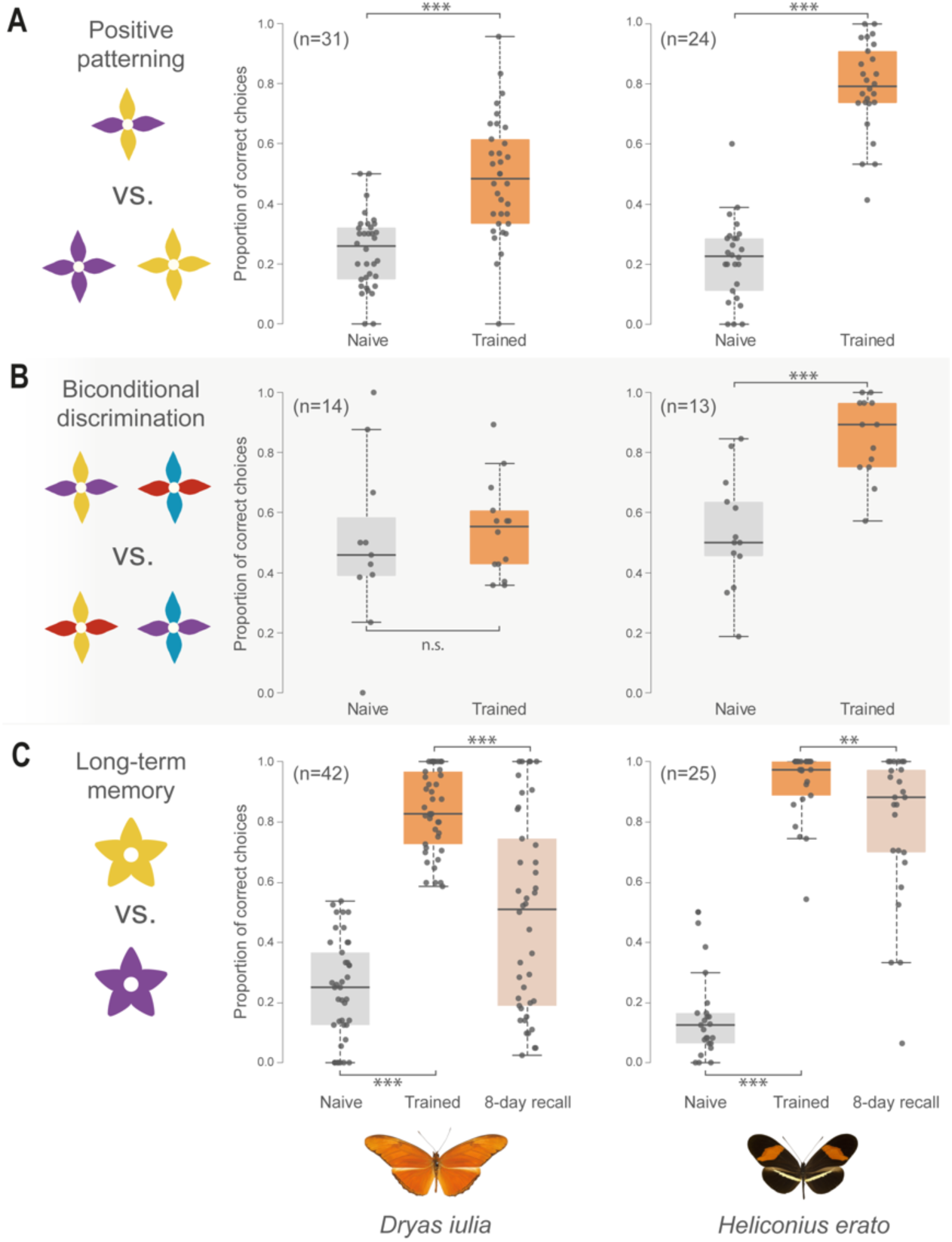
Enhanced non-elemental visual learning and long-term memory in *H. erato* relative to *Dryas iulia*. (**A**) Both species solve a visual positive patterning task, but *H. erato* is significantly more accurate. (**B**) *H. erato* can solve the more difficult and strictly non-elemental biconditional discrimination task, whereas *Dryas iulia* cannot. (**C**) *H. erato* has superior memory of a learned visual cue after 8 days compared with *Dryas iulia*. * = p < 0.05; ** = p < 0.01; *** = p < 0.001.

Trap-line foraging also requires the formation of stable long-term memories to exploit resources repeatedly over many weeks or months (*14*). Mushroom bodies are central to long-term olfactory memories in insects (*49, 50*), and while their role in long-term visual memory is less explored, it is likely conserved across modalities. To test whether enlarged mushroom bodies may be associated with improved memory retention, we again trained *D. iulia* and *H. erato* on artificial feeders, this time in a simple two-color preference assay (purple/yellow). After demonstrating that both *H. erato (*Z ratio = −12.136, p < 0.001) and *Dryas iulia* (Z ratio = −11.321, p < 0.001) learn to associate a reward with a non-preferred color cue (Figure 5C), we fed the butterflies with neutral (white) feeders for eight days. In subsequent preference trials, *D. iulia* performed significantly worse than *H. erato* (Z ratio = −4.829, p < 0.0001), which maintains a high accuracy towards the reinforced cue (Z ratio = 4.635, p < 0.0001), while *D. iulia*’s preference is not significantly different to random (Z ratio = −0.160, p = 0.873) (Figure 5C). While further exploration of the cognitive shifts supported by mushroom body expansion are warranted, our data provide evidence that *Heliconius* have improved configurational learning, and more accurate long-term memory. These two functions are likely essential to meet the cognitive demands of learning the location of resources across space and time.

## vii. Conclusions

By combining extensive, phylogenetically dense sampling of wild individuals, state-of-the-art comparative analyses, and focused behavioral experiments, our data demonstrate considerable variation in a prominent brain structure within a closely related and - with the exception of the dietary innovation of pollen feeding - ecologically comparable tribe of Neotropical butterflies. Notably, the expansions we observe in this tribe, which diverged across the last ~25mya, are an order or magnitude more recent than other major mushroom body expansion events (*51*), meaning comparative studies suffer from fewer confounding effects. Taking this approach, we provide evidence of mushroom body expansion associated with the requirements for more refined visual processing, supported by a massive increase in Kenyon cell number. While our analyses suggest the ecological factors which favor increased investment in mushroom bodies across Heliconiini may be more complex than previously hypothesized, critically, we provide evidence that it is accompanied by enhancements in visual learning and memory, with the most substantial shifts in size and sensory modality co-occurring with the origin of pollen feeding and trap-line behavior. Our data therefore highlight Heliconiini as a tractable system for comparative, detail-rich analyses aimed at understanding the adaptive and mechanistic basis of neural and behavioral evolution.

## Acknowledgements

We are very grateful to the environmental ministries of Costa Rica, Panama, French Guiana, Ecuador and Peru for permission to collect and export samples. We thank the Organization for Tropical Studies, Le Leona Eco Lodge, the Smithsonian Tropical Research Institute, the Estación Científica Yasuní, PUCE, F. Ramirez Castro, Neil Rosser, Ronald Mori Pezo the Dasmahapatra group, and the broader *Heliconius* research community for support in the field and for discussions, and to Swidbert Ott for advice and encouragement early on in this project. We are grateful to the Wolfson Bioimaging Centre, University of Bristol, the University College London Confocal Imaging facility, and Matt Wayland and the Dept. of Zoology Imaging Facility, University of Cambridge, for imaging assistance.

## Funding

Royal Commission for the Great Exhibition Research Fellowship (SHM).

Leverhulme Trust Early Career Fellowship (SHM).

Short-term STRI Fellowship (SHM).

Royal Society Research Grant (SHM).

ERC Starter Grant 758508 (SHM).

NERC Independent Research Fellowship NE/N014936/1 (SHM).

PhD Studentship from Trinity College, Cambridge (FJY).

## Author contributions

Conceptualization: SHM

Methodology: SHM, AC, FJY, CN, FC

Investigation: AC, FJY, DA, SM, LMF, LH, MM, CN, FC

Visualization: FJY, AC

Funding acquisition: SHM, WOM

Project administration: SHM, WOM

Supervision: SHM, WOM

Writing – original draft: SHM, FJY, AC

Writing – review & editing: SHM, AC, FJY, WOM, FC, CN, DA, SM, LMF, LH, MM.

## Supplementary Materials

### 1. Supplementary methods

#### 1.1 Phylogenetic sampling across Heliconiini

Heliconiini are distributed across Central and South America and occur in a range of habitats from dry to wet forest, and from sea level to ~2500m. To ensure a thorough and even sampling across the phylogeny and avoid biased phylogenetic analyses, a series of sampling trips were performed to collect wild butterflies across the tribe. We are indebted to the environmental agencies in Costa Rica, Panama, French Guiana, Ecuador and Peru for permissions to carry out this work. Sampling of wild individuals (Table S1) was focused on the following localities:

**i) Costa Rica:** Sampling was performed in 2015 at La Selva Biological Station (elevation 30-130m), Las Cruces Biological Station (elevation <20m), Le Leona eco-lodge on the edge of Corcovado National Park (elevation ~1000m), and Orosí (elevation ~1300m). Samples were collected under permit SINAC-SE-GASP-PI-R-2015. We thank the Organization for Tropical Studies at Las Cruces and La Selva, and Le Leona Eco Lodge for assistance in Costa Rica for support and assistance.
**ii) Panama:** Sampling was performed in 2012 and 2013 along Pipeline Road, Gamboa (elevation 60 m), which transects open to closed forest, and the nearby Soberanía National Park. Samples were collected under permits SEX/A-3-12, SE/A-7-13 and SE/AP-14-18. We are grateful to the *Heliconius* community at the Smithsonian Tropical Research Institute for valuable support during this period.
**iii) French Guiana:** Sampling was performed in 2016 at several urban sites round Cayenne and along roadsides (N1, N2, D5, D6) in forest edge habitats in the Arrondissement of Cayenne (elevation 0-150m). At the time of sampling no permits were required to sample outside National Parks in French Guiana.
**iv) Ecuador:** Sampling was performed at the Estación Científica Yasuní, in the Parque Nacional Yasuní, Orellana Province, Ecuador in November/December 2011 and September/October 2012 under permit collection no. 0033-FAU-MAE-DPO-PNY and exported under permit nos. 001-FAU-MAE-DPO-PNY and 006-EXP-CIEN-FAU-DPO-PNY. These were obtained from Parque Nacional Yasuní, Ministerio Del Ambiente, La Dirección Provincial de Orellana with the help of Estación Científica Yasuní and Pontificia Universidad Católica del Ecuador (PUCE). We are thankful to Á. Barragán, E. Moreno, P. Jarrín, and D. Lasso from the Estación Científica Yasuní, Pontificia Universidad Católica del Ecuador, and M. Arévalo from the Parque Nacional Yasuní Ministerio del Ambiente for assisting with collection and export permits. We are also grateful to F. Ramirez Castro for his assistance in the field in 2011. Additional collections were made from the forests around Vilcabamba (~1200m) and Balsas Canton (~500m) in Southern Ecuador in 2012 under permits provided by Ministerio del Ambiente del Ecuador to Dr Richard Merrill.
**v) Peru:** Sampling was performed in 2014 around the Escalera region near Tarapoto, Departamento de San Martín (elevation 300-1295 m), around Schucshuyacu (0-50m) on the Huallaga River in lowland forest to the east of Tarapoto, and along several higer eleveation sites to the northwest of Tarapoto: Abra Patricia at the San Martin/Amazonia boder (2350m), Catarata Yubilla, Cuispes (2100m), Catarata Gocta, Cocachimba (2300m), and roadside sites near Pedro Ruiz Gallo (1300m). Additional sampling was performed in Excalera in 2015. Samples were collected under permits 0289-2014-MINAGRI-DGFFS/DGEFFS, 020-014/GRSM/PEHCBM/DMA/ACR-CE, 040–2015/GRSM/PEHCBM/DMA/ACR-CE, granted to Dr Neil Rosser. We thank Neil Rosser, Ronald Mori Pezo and the Dasmahapatra group for assistance in Peru.

All individuals were collected using hand nets and kept alive in glassine envelopes until brain tissue could be fixed within a few hours of collection. Dissections (see below) were carried out at accommodation sites in the field. Body mass, length and wingspan were recorded as measures of body size and wings were kept in glassine envelopes as voucher specimens.

In addition to the wild caught samples, we obtained additional samples for selected species representing key lineages for subsequent analysis of the cellular composition and sensory domains of the calyx using pupae obtained from commercial supplies (Stratford Butterfly Farm, UK/London Pupae Supplies, UK/ Costa Rica Entomological Supplies, Costa Rica). Once dry, freshly eclosed butterflies were individually labelled and aged in a 2m^3^ cage (2m × 1m × 1m) in a controlled temperature room at 28°C with 80% humidity and a 12-hour day/night cycle. The cage was enriched with two plants, *Passiflora biflora* and *Lantana sp*., a natural food resource, with supplemental feeding using artificial feeders containing 30% sugar solution with ~10% bee collected pollen. For the neural tracing experiment and for synapse/microglomeruli quantification, the butterflies were sampled six to ten days after emergence, at which point they are considered behaviourally and sexually mature (Mallet 1981). For the quantification of Kenyon cells adult butterflies were sampled at least 24 hours after eclosion. as pilot data suggested major differences in mushroom body size are apparent at eclosion, suggesting the major shift in Kenyon cell production is completed by the adult stage.

#### 1.2 Fixation

For all samples, following any additional procedures (see below), brains were dissected out in isotonic buffer-saline (HBS: 150 mM NaCl, 5 mM KCl, 5 mM CaCl2, 25 mM sucrose, 10 mM HEPES, pH 7.4) and fixed for 16 to 20 hours in zinc-formalin solution (ZnFA; 18.4 mM ZnCl2, 135 mM NaCl, 35 mM sucrose, 1% formalin), under gentle agitation at room temperature. The brain was then washed (3x 10 mins in HBS) and subsequently dehydrated into a solution containing 80% methanol and 20% dimethylsulfoxide (DMSO) for 2 hours, before storage in 100% methanol at −20°C. When ready to process, samples were brought back at room temperature and rehydrated in bath series of decreasing methanol concentration (90%, 70%, 50%, 30%, 0% in 0.1 M Tris buffer, pH 7.4, for 10 mins each).

#### 1.3 Neuropil staining

For neuropil staining we followed Ott’s (1) protocol. Mono-clonal anti-synapsin antibody (mouse anti-SYNORF1: 3C11, DSHB, RRID:AB_2315424), which bind a vesicle-associated protein at presynaptic site, was used to reveal the neuropil structure of the brain in all samples. To prevent antibodies from unspecific binding, brains were first incubated for 2 hours in a blocking solution (NGS-PBSd) containing 5% of normal goat serum (NGS: G9023, Sigma-Aldrich) in 0.1 M phosphate-buffered saline with 1% DMSO (PBSd). The primary antibody 3C11 (anti-SYNORF1) was then applied at a 1:30 dilution, in PBSd-NGS for 3.5 days at 4°C under agitation. Non-bound antibodies were rinsed out using three consecutive baths of PBSd solution (2h each) before incubation with Cy2-conjugated secondary antibody (Cy2 goat anti-mouse IgG: 115-225-146, Jackson ImmunoResearch, RRID: AB_2307343) diluted at 1:100 in PBSd-NGS for 2.5 days at 4°C. The brains were then dehydrated in bath series of increasing glycerol concentration (1%, 2%, 4% for 2 hours each, and 8%, 15%, 30%, 50%, 60%, 70%, 80% for 1 hour each) in 0.1 M Tris buffer with 1% DMSO, followed by 3 washes in 100% ethanol (30 mins each). The brain tissues were clarified in methyl salicylate (M6752, Sigma-Aldrich) for 30 mins before being transferred in fresh methyl salicylate, which was used as storage and mounting medium.

#### 1.4 Neuronal tracing

To measure the volume of visual and olfactory areas within the mushroom bodies, projection neurons from primary sensory neuropils (the antennal lobe and the optic lobe) were differentially labelled using fluorescent tracers. Butterflies were immobilized in custom-made holders with the head isolated from the rest of the body using a plastic collar placed at the neck. To prevent the solution from leaking, a waterproof area was built by applying beeswax, melted at low temperature, around the head. The head capsule was then opened and the brain was exposed under a ringer solution (150 mM NaCL, 3mM CaCl_2_, 3 mM KCL, 2 mM MgCl_2_, 10 mM Hepes, 5 mM glucose, 20 mM sucrose) so that the tracheal sac and glands above the sensory neuropils could be removed. Under filtered illumination, the tip of pulled glass capillaries (G100-4, Warner Instruments) was loaded with crystals of dextran-conjugated dyes; either dextran-tetramethylrhodamine (fluoro-ruby: 10,000 MW, D1817, Thermo Fisher Scientific) or dextran-Alexa fluor 647 (10,000 MW, D22914, Thermo Fisher Scientific) mixed in 2% Bovine serum albumin (BSA). The loaded tip was either injected into the target neuropil(s), initially the mushroom body calyx, then subsequently the antennal lobe or in the caudal junction between optic lobe and the central brain (aiming for output neurons of the ventral lobe of the lobula), which were identified as major input sites to the calyx (Figure S8, see below). The glass capillary was kept in this position until the dye crystal dissolved so that neurons were imbued with the fluorescent tracer. After each injection, extra dye was removed by repeatedly washing the brain with Ringer solution, and absorbing the liquid with thin paper wipes. The heads were then covered with fresh Ringer solution and the butterflies were kept in the holders overnight in the dark. The next day, the brains were dissected out and preserved (as described in 1.2), for later background staining of the neuropils (as described in 1.3).

#### 1.5 Sectioning and mounting

To visualize the cell bodies of Kenyon cells and calycal synapse density at high magnification, the butterfly brains were physically sectioned to overcome limitations of the working distance of high magnification lenses. Preserved samples were rehydrated (as described in 1.2) and embedded in 5% low melting point agarose (16520-050, Thermo Fisher Scientific). The brains were sliced horizontally along the anterior-posterior axis, in frontal sections with a thickness of 80 µm using a vibratome (VT 1000 S, Leica Biosystems, Wetzlar, Germany). The sections were collected and stained following 2 different procedures (described in 1.6 for Kenyon cells or in 1.7 for synapses/microglomeruli). After the staining procedure, brain sections were kept in 60% glycerol overnight, then mounted on microscope slides in 80% glycerol and surmounted by cover slips which were sealed with commercial nail polish.

#### 1.6 Kenyon cell staining

To identify Kenyon cells over non-neuronal cells, cell nuclei were stained in combination with a specific marker of neuronal membrane, the anti-peroxidase antibody (HRP: Rabbit anti-horseradish peroxidae, P-7899, Sigma-Aldrich, RRID:AB_261181). Brain slices were first permeabilized in PBS containing 2% of Triton X-100, for 30 mins at room temperature. Slices were then blocked in PBS-NGS (PBS with 5% normal goat serum) with 0.2% Triton (PBSt-NGS thereafter). The brain tissues were probed with rabbit anti-HRP applied at a dilution of 1:10 000, in PBSt-NGS for 3 days at 4°C, under a constant agitation. Next, the slices were washed in PBSt (3 x 30 minutes), to remove residual unbound antibodies, and subsequently stained with Cy3-conjugated secondary antibody (Cy3 goat anti-rabbit IgG: 111-165-144, Jackson ImmunoResearch, RRID: AB_2338006), diluted at 1:200 in PBSt-NGS. After 3 days of incubation at 4°C, the brain sections were washed in PBSt (3x 30 mins) and stained with 1:1000 DAPI (D9542, Sigma-Aldrich) for 30 mins at room temperature. Extra reagent was then washed out and the brain slices were mounted on microscope slides (as described in 1.5).

#### 1.7 Synapse/Microglomeruli staining

To assess the presynaptic boutons density within the calyx, brain slices were stained using anti-SYNORF1 antibody, or a combination of anti-SYNORF1 and Alexa-conjugated phalloidin which selectively label F-actin filament allowing to reveal dendritic profiles of Kenyon cells. For six species (Figure 3D) we focused solely on synaptic density using automated synapse counting of ZnFA fixed, anti-SYNORF1 stained sections (see 1.9), but we additionally stained using both primary antibodies for two species (Figure 3E; *H. charithonia* and *D. iulia*) to manually count microglomeruli, synaptic complexes containing presynaptic boutons and surrounding post synaptic Kenyon cell dendrites (2). As methanol destroys the quaternary structure of actin proteins, phalloidin stained samples were first fixed in a solution of methanol-free formaldehyde (18.4 mM ZnCl2, 135 mM NaCl, 35 mM sucrose, 1% formaldehyde) to preserve its native conformation. After fixation, both sets of samples were sliced (as described in 1.5) and stained as follows. Brain slices were permeabilized in PBS containing 2% Triton for 10 mins, and subsequently washed in PBSt (2x 10 mins). The samples were then blocked for 2 hours in PBSt-NGS and probed with anti-SYNORF1 antibody (3C11) at 1:30 in PBSt-NGS for 4 days at 4°C. After 3 consecutive washes in PBSt (3x 2h), the slices were stained in Alexa 568-conjugated secondary antibody (Alexa 568 goat anti-mouse IgG: A-11004, Thermo Fisher Scientific, AB_2534072) diluted at 1:200, and Alexa 488 phalloidin (A12379, Thermo Fisher Scientific) diluted at 0.5 units/mL in PBSt-NGS, for 7 days at 4°C. Finally, the slices were washed in PBSt (3x 20 mins) and mounted on microscope slides (as described in 1.5).

#### 1.8 Confocal image acquisition

For whole mount brains used to quantify neuropil volumes, clarified samples (see 1.3) were mounted in wells drilled into custom made 3mm thick aluminium-slides, filled with methyl salicylate, and sealed with cover slips on both sides. Whole-mount brain imaging was performed using confocal laser-scanning microscopes (Leica SP5 or SP8, Leica microsystems, Wetzlar, Germany) using a 10X dry objective with 0.4 NA (10x HC PL APO CS, Leica microsystems No.11506285), a mechanical z-step of 2µm, and an x-y resolution of 512 x 512 pixels. For most species, imaging the whole brain required capturing 2 × 2 or 3 × 2 tiled stacks in the *x-y* dimensions (with 20% overlap) that were automatically merged in Leica Applications Suite Advanced Fluorescence software. In addition, where necessary brains were scanned from the posterior and anterior side to span the full *z*-dimension of the brain, due to low image quality with increasing *z*-distance. These two image stacks were then merged in Amira 3D analysis software 5.5 (FEI Visualization Sciences Group; custom module ‘Advanced Merge’). Images of visual and olfactory projections in the MB (after 1.4) were obtained using the same microscopes and objective lens. The MB calyxes were scanned from the ventral surface of the brain at a resolution of 1024 x 1024 pixels (*x*,*y*) every 2µm (z). The “SYNORF1-cy2” neuropil staining was excited with an argon laser at 488nm wavelength, whereas traced projections with dextran-conjugated dyes (Tetramethylrhodamine and Alexa fluor 647) were excited at 561nm and 633nm respectively, with solid state lasers. To avoid crosstalk, each wavelength was sequentially scanned between lines, and emitted light was received on different detectors, a hybrid detector (Leica HyD, for cy2) and two standard PMTs. The *z*-dimension was scaled 1.52× to correct the artefactual shortening associated with the 10× air objective.

Images of brain slices (after 1.5-1.7) were obtained using the same confocal microscopes. To acquire images of the MB calyx and all surrounding Kenyon cell clusters, every brain section stained with “HRP-cy3” (after 1.6) was scanned at a resolution of 1024 x 1024 pixels (*x*,*y*) every 3µm (z) with a 10X dry objective (0.4 NA) and a solid state laser excitation at 561nm wavelength. Subsamples of the Kenyon cells cluster were then scanned at higher magnification with a 63X glycerol immersion objective with a 1.3 NA (63X HCX PL APO CS, Leica microsystems No. 11506194), and UV excitation at 405 nm wavelength. Images stacks measuring about 150 x 150 x 30 µm were produced for five randomly selected areas within the Kenyon cell clusters for each brain, at a resolution of 1024 x 1024 pixels (*x*,*y*) and a 1µm *z*-ztep. Images of presynaptic boutons stained with anti-SYNORF1 antibodies, and microglomeruli stained with Alexa-conjugated phalloidin (after 1.7) were acquired in a similar manner, using five randomly selected areas of each calyx with the same 63X glycerol objective. Boxes of approximately 125 x 125 x 25 µm were scanned to measure the density of presynaptic boutons, and 95 x 95 x 25 µm for the microglomeruli density estimates, both at a resolution of 1024×1024 pixels every 1 µm along the depth (z).

For these analyses, the average densities from these subsampled regions were multiplied by the total volume or the Kenyon cell cluster/calyx to produce total numbers. Since the brain slices mounted in glycerol under cover slips were scanned with a 10X dry objective, there is an axial shift of the emitted light through different optical media resulting in mismatching refractive indices. To rescale images of brain sections, a correction factor was therefore assessed by scanning calibration beads (FocalCheck Microspheres, 15 µm, fluorescent green/orange/dark-red ring stains, F7235, Thermo Fisher Scientific) and calculating the ratio between measured z (depth) size and the actual size (15 µm). The average ratio obtained over 5 measurements of the microspheres (1.85) was used as correction factor of the z-dimension.

#### 1.9 Confocal image analysis

For 3D segmentations from whole brain samples, confocal image stacks were saved as .lif files using the Leica application suite (RRID:SCR_013673) and imported in Advanced 3D Visualization and Volume Modelling software (Amira 5.4.3; Thermo Fisher Scientific, RRID:SCR_007353) to build virtual models of the neuropils. The z-dimension was rescaled by 1.52 when the mounting medium was methyl salicylate, whereas a correction of 1.85 was applied for brain slices scanned in glycerol (as described in 1.8). Image regions were assigned to anatomical structures with the *labelfield* module of the software, by defining separate outlines based on the brightness and contrast obtained in the different light channels (synaspsin: 488nm, neural tracing: 561 and 633 nm, HRP: 561nm). For large neuropil, approximately every fourth image was manually segmented and interpolated in the z-dimension across all images that contain the neuropil of interest, for smaller neuropils images were labelled at higher frequency. The volume of each 3D neuropil model was then extracted using the material statistics module of the Amira software. In total, the brains of 318 wild-caught individuals, representing 41 Heliconiini species, were reconstructed. For each of these brains, we reconstructed the volume of the mushroom body calyx, peduncle and lobes, two optic lobe neuropils, the medulla and ventral lobula, and the antennal lobe from one hemisphere, multiplying by two to get the total volume across the brain. In addition, we measured the total volume of the central brain (CBR), then subtracted the total volume of the mushroom body and antennal lobes to get an independent measure of brain size as an allometric control (rest-of-CBR, rCBR). For neural tracing experiment, the mushroom bodies and its sensory areas were segmented in 63 individuals belonging to 8 species (8 individuals/species with the exception of *H. doris* [7 individuals]) reared in controlled conditions, and for the Kenyon cell/synapse counting experiment the calyx and cell cluster of 50 individuals belonging to 6 species (approximately 8 individuals/species with the exception of *A. vanillae* [10 individuals for Kenyon cells]) were reconstructed from sectioned brains.

To assess the number of Kenyon cells across Heliconiini butterflies, the density of cell bodies was measured by generating cubes of 25 x 25 x 15 µm from each high magnification image stack of Kenyon cells clusters (5 per individual; described in 1.8) using ImageJ software (RRID:SCR_003070) and the Bio-Formats library (RRID:SCR_000450). The number of nuclei within each cube was manually counted, omitting any cells with cell membranes not stained with HRP (see 1.6), and the average density was then multiplied by the volume of the whole Kenyon cells cluster around the MB (measured as described above) to obtain an estimated number of Kenyon cells per hemisphere for each individual.

The synaptic density within the calyx was measured from staining of presynaptic boutons with the anti-SYNORF1 antibody, using the ImageJ plugin 3D Object Counter (RRID:SCR_017066). The number of synapses was automatically counted within five boxes of 50 x 50 x 15 μm, adjusting intensity threshold to each image stack. Background noise was filtered by excluding all detected signals smaller than 10 voxels from the count, as manual measurements of synapse size suggested objects below this are unlikely to be genuine synapses. The density of microglomeruli in *Heliconius charithonia* and *Dryas iulia* was measured in a similar way as described for the Kenyon cells (above) in five of 25 x 25 x 10 µm cubes for each individual, generated from confocal image stacks of Alexa-conjugated phalloidin staining.

#### 1.10 Electron microscopy sample preparation and imaging

To confirm cell count methods using immunohistochemistry and imaging of the Kenyon cell cluster, we obtained additional estimates using Electron Microscopy to image a cross section of the peduncle, to visualise Kenyon cell axons. For this purpose, butterflies were stunned/anaesthetised with Carbon dioxide prior to immersion in 2.5% glutaraldehyde in 0.1M cacodylate buffer at room temperature. Samples were then stored in a 4C fridge. Brains were subsequently removed, fixed and dehydrated using standard techniques. Briefly, treatments were made with osmium tetroxide (2% in cacodylate buffer), enbloc stained with uranyl acetate, dehydrated with ethanol, and infiltrated with propylene oxide, and then Epon resin mixture. Embedded blocks were polymerised at 60°C prior to sectioning on a Reichert Ultracut E. Transverse sections were taken of the peduncle to reveal axonal structure and imaged using a Tecnai T12 electron microscope (ThermoFisher UK) at 120 kV.

#### 1.11 Ecological data

We scored three ecological traits for each Heliconiini species: the presence of pollen feeding, the number of host plant species exploited, and the degree of social roosting. We categorised pollen feeding as a binary trait, which was true for all *Heliconius,* except for *H. aoede,* and false for all other Heliconiini (3). Host plant use number was taken from the dataset collated by Kozak (4). Roosting behaviour was classed as either solitary, loose group, small group, or large group, taken from Brown (5).

#### 1.12 Phylogenetic trees

All phylogenetic comparative analyses were conducted using a new phylogenetic tree of the Heliconiini generated from newly assembled genomes (6). To assemble additional genomes of *Heliconius erato cyrbia*, *H. melpomene rosina*, *H. m. amaryllis*, and *H. cydno galanthus*, short-read illumina data was downloaded from NCBI (National Centre for Biotechnology Information) and a reference-guided assembly approach adapted and extended from Lischer and Shimizu (7) adopted. The strategy involves first mapping reads against a reference genome of a related species (*H. e. demophoon* for *H. e. cyrbia*; *H. m. melpomene* 2.5 for *H. m. rosina*, *H. m. amaryllis*; *H. cydno* for *H. cydno galanthus*) to reduce the complexity of *de novo* assembly within continuous covered regions, then later integrating reads with no similarity to the related genome.

Extra scaffolding procedures were implemented in order to improve the previous reference guided *de novo* assembly pipeline (7). Leveraging the very small genetic distances of these subspecies with their reference genomes, RNA-seq data from the reference species were downloaded from NCBI concatenated, corrected a normalized using *BBMap* v 38.79 (8) [target=20 maxdepth=20 mindepth=5]. These reads were mapped using *HISAT2* v 2.1.0 (9), and *P_RNA_scoffolder* (10). Following this step, *RaGOO* (11) was used [-T sr] for homology-based scaffolding and misassembly correction. *RaGOO* identifies structural variants and sequencing gaps, and accurately orders and orients *de novo* genome assemblies. *abyss*-*Sealer* (12) was then used with multiple *kmer*s [-k99 -k97 -k95 -k93 -k91 -k89 -k85 -k81 -k77 -k73 -k69 -k65 -k61 -k57] to finalise the assembly to close remaining gaps. After the genome assembly was completed, contaminants were identified using *BlobTools* v 1.1.1 (13) and removed from the final assemblies.

Single-copy orthologous genes in each genome were then identified using *BUSCO* (Benchmarking Universal Single-Copy Orthologs; v3.1.0 (14)). The Insecta set was selected in *OrthoDB* v 9 (odb9 1958 genes) using default parameters [-m genome]. For genes recovered as fragmented and missing a further mapping was implemented with *Exonerate*, using protein sequences found in other *Heliconius* species to deal with possible false discovery hits. For each species, all *BUSCO* genes found in a single copy were used for phylogenetic analysis.

Each nucleotide sequence *BUSCO* locus was aligned separately with *MACSE* v 2 (15), and all alignments were concatenated. From the concatenated alignment, gaps were removed using *Gblock* v 0.91b (16) [−t=c − b1=(#Nseq/2+1) −b2=(#Nseq/2+1) −b3=1 −b4=6 −b5=h] following Cicconardi et al. (2020; 2017). A maximum likelihood (ML) phylogenetic tree was performed using *IQ-TREE* v 2 (19), partitioning the supermatrix for each locus and codon position. *IQ-TREE* was run with the following settings: --runs 5 -m MFP. 5,000 ultrafast bootstrap replicates were conducted, resampling partitions and then sites within resampled partitions (20, 21) to reduce false positives (-b 5000 -- sampling GENESITE).

The Bayesian algorithm of *MCMCTree* (22) was implemented with approximate likelihood computation to estimate divergence times. First, branch lengths were estimated by ML, and then the gradient and Hessian matrix around these ML estimates was calculated in *MCMCTree* using the DNA supermatrix. Calibration nodes were constrained according to Cicconardi et al. (in prep). using a uniform distribution. To ensure convergence, the analysis was run for 10 x 100k generations after a 10M generations as burn-in, logging every 200 generations. *Tracer* v 1.7.1 (23) was used to check for convergence and ESS values were > 200.

### 2. Statistical analysis

#### 2.1 Volumetric comparisons across Heliconiini

##### i) GLMs and allometric scaling

All volumetric measurements were log_10_-transformed for statistical analyses. Where possible, analyses were conducted using individual-level data, though certain analyses required the use of species averages, calculated as the arithmetic mean. All analyses of wild data were run using the Helicoiniini phylogeny generated as described in 1.12, and were performed in R v 4.1.2. We ran a series of phylogenetic generalised linear mixed models (GLMM) with gaussian distributions using the R package *MCMCglmm* v 2.32 (24) to determine whether the scaling relationship between the mushroom body and the rest of the central brain (rCBR) changes between *Heliconius* and non-*Heliconius* Heliconiini, and between Heliconiini genera and subclades within *Heliconius* (25), including the effects of body size measurements and sex, where appropriate. We also included antennal lobe (AL) and medulla (ME) volumes to test whether increases in sensory neuropils, and therefore possibly increased sensory inputs, could provide a simple explanation of variation in mushroom body size. The best fitting model was then identified using Deviance Information Criterion (26). To determine whether any increases in MB size are driven by the expansion in a particular region of the mushroom body, we tested for variation in the scaling relationship between the mushroom body calyx and the lobe and peduncle between *Heliconius* and non-*Heliconius* species. We also assessed whether the mushroom bodies are unique in exhibiting expansion in *Heliconius* by testing whether the genus shows an increase in medulla, ventral lobula (vLO) or antennal lobe size, controlling for rCBR. Additionally, we tested for variation in rCBR size, using body size measurements as an allometric control. All models were checked for convergence using the *gelman.diag* function and for auto-correlation using the *autocorr* function provided in *MCMCglmm*, in addition to visually inspecting the trace plots. All models were run for 500,000 iterations, with a burn-in of 10,000 and a thinning factor of 500.

To support these analyses, we performed an additional set of analyses to examine allometric shifts between mushroom body size and rCBR. First, the function *sma* from the R package *smatr* v 3.4-8 was used to test for pairwise interspecific differences in the scaling relationship between mushroom body volume and rCBR across all species, testing for both differences in the slope and elevation of scaling relationships (27). For this analysis, only species with at least 8 individuals were included, reducing the dataset to 26 species, and the “robust” option was set to true for these analyses to minimise the influence of potential outliers (28).

##### ii) phylogenetic modelling using BAYOU

The R package *bayou* v 2.0 was used to identify regions of the Heliconiini tree showing evidence of a shift in the scaling relationship between mushroom volume and rCBR without *a priori* constraints (29). This method fits multi-optima Ornstein-Uhlenbeck models to phylogenetic comparative data, estimating the placement and magnitude of adaptive shifts. We compared three models: one with no shifts in either slope or elevation (all species are assumed exhibit the same allometric scaling relationship), one allowing for shifts in elevation (a common allometric slope is assumed, but “grade-shifts” are allowed), and one allowing for shifts in both elevation and slope. Models were each run for 1,000,000 iterations, with a burn-in of 300,000 generations and their fits were compared using the *steppingstone* function in *bayou*, run for 10,000 generations with a burn-in of 3000 (30). The posterior probability cut-off for identifying a shift in the relationship between mushroom body and rCBR size was set at 0.5 following published recommendations (29,31,32).

##### iii) analysis of evolutionary rates and ancestral state reconstruction

To identify key periods of evolutionary change in a robust fashion, we used two different methods to test for shifts in the evolutionary rate of change in mushroom size within the Heliconiini, checking for consistency in the interpretations implied from the output of each method. First, we used *BayesTraits* v 3 to compare two independent contrast MCMC models of evolution, one allowing for a rate scaling parameter to vary across branches, and a second one where it is averaged across all branches (33, 34). In the variable rates model, branch lengths are scaled to accommodate periods of increased trait change, and this scaling parameter provides an indication of evolutionary rate. Models were run for 110,000,000 iterations, with a burn-in of 10,000,000, and sampled every 10,000 iterations. Model fit was compared by calculating Log Bayes Factors using marginal likelihoods, calculated for each model using the stepping stone sampler (35) and sampling 100 stones for 10,000 iterations each, Log Bayes Factors are calculated as two times the difference in marginal log likelihoods of the two models, where a Log Bayes Factor <2 is interpreted as weak evidence for the more complex model, >2 as positive evidence, 5-10 as strong evidence, and >10 as very strong evidence. The output from the variable rates model was processed using the online tool (www.evolution.reading.ac.uk/BayesTraitsV4.0.0/BayesTraitsV4.0.0.html), to determine scaling factors for each node.

Second, we used a recently-published method that uses Brownian motion to model variations in evolutionary rate (36). Under this model, different branches of the tree are assumed to have different evolutionary rates, which also evolve via a Brownian process. Unlike the Bayesian approach implemented in *BayesTraits*, this method involves continuous (rather than discrete) changes in evolutionary rate between, or along, branches. This method has been implemented in the *multirateBM* function in the R package *phytools* v 0.7-90 (37). We used this method to estimate variations in the evolutionary rate of both mushroom body size. We then calculated the residual evolutionary rate of mushroom body size for each branch through a linear regression, using the *lm* function in R, of the estimated evolutionary rates of the mushroom body and rCBR size.

For these analyses of evolutionary rates, we used an expanded dataset that also included eight outgroup Lepidoptera species to better estimate the ancestral state at the *Heliconiini* root node. Neuropil volumes for the outgroup species were collected from published data (38–43), with the exception of *Bicyclus anynana* which was newly collected by S. Montgomery. Divergence times for outgroup species were taken from a recently published Lepidoptera phylogeny (44).

Finally, we also used two methods to estimate ancestral states for mushroom size and rCBR size at key internal nodes within the Heliconiini tree. First using *BayesTraits* v 3 we compared a random walk and directional MCMC model of evolution in mushroom body size, controlling for rCBR, using the series of scaled trees generated from the post-burnin iterations of the variable rates model described above. The negative Log Bayes Factor (−16.84) for these two models indicated that a directional model of evolution was not supported. we then used the non-directional model and the scaled trees to estimate values for MB size and rCBR size at key internal nodes. Second, we used the *fastAnc* function in the *phytools* package to estimate the maximum likelihood ancestral states for mushroom body size and rCBR size at each node in the Heliconiini tree. The ancestral state density plots in Figure 2B we generated using the distributions of ancestral states estimated using *BayesTraits* v 3.

##### iii) Re-analysis of Snell-Rood et al. (2020) dataset

To corroborate our findings, we re-analysed a published, independently collected, dataset of neuroanatomy for 41 species of North American butterfly, which included *H. charithonia* and the Heliconiini *Agraulis vanillae*. Using this dataset, we ran the BAYOU analysis (2.1.ii), and the evolutionary rates analyses (2.1.iii) as described above. We used a recently published phylogeny of North American butterflies to perform these analyses (45).

#### 2.2 Variation in Kenyon cell number, calyx synapses and microglomeruli

All count data were log transformed to meet the assumptions of being normal distributed. We tested for interspecific variation in Kenyon cell number, and separately Kenyon cell density, with generalised linear models (GLMs) with species as a fixed effect. We tested for variation in total Kenyon cell number, and separately Kenyon cell density, between *Heliconius* and non-*Heliconius* individuals using generalised linear mixed models (GLMMs) with a gaussian distribution, with membership in the *Heliconius* genus as a fixed effect and species as a random effect. We then explored whether the relationship between Kenyon cell number and calyx volume varies between taxa using a GLM with Kenyon cell number and species as fixed effects, and whether it varies between *Heliconius* and non-*Heliconius* using a GLMM with *Heliconius* membership as fixed effects and species as a random effect. A similar approach was adopted for synaptic data: interspecific differences between synapse density and total synapse number were examined separately using GLMs with species as a fixed effect. Differences between *Heliconius* and non-*Heliconius* individuals were also tested using GLMMs with a Gaussian distribution and *Heliconius* membership as a fixed effect and species as a random effect. Finally, variation in microglomeruli density between *Heliconius charithonia* and *Dryas iulia* was tested using a GLM with species as a fixed effect.

All models were built in R using the *glm* function for GLMs and the *glmer* function from package *lme4* v 1.1-30 for GLMMs. Diagnostics were assessed using the package *DHARMa* v 0.4.4 for R (46). All pairwise differences were assessed by calculating the estimated marginal means using the function *emmeans* in the R package *emmeans* v1.7.0, correcting for multiple comparisons using the Tukey test (47). We also independently assessed whether the relationship between Kenyon cell number and calyx volume varies between species using the function *sma* from the R package *smatr* v 3.4-8. We tested for pairwise differences between species in both the slope and elevation of the scaling relationship between these traits. The “robust” option was set to true for these analyses to minimise the influence of potential outliers (28).

#### 2.3 Variation in sensory input into the calyx

Visual and olfactory calyx volumes were log transformed for better normal distribution. Overall differences between *Heliconius* and non-*Heliconius* individuals in visual and olfactory calyx volume were analysed separately using GLMMs with *Heliconius* membership as a fixed factor and species as a random factor. We then determined whether the variation was higher in the visual or olfactory calyx by comparing the effect sizes of *Heliconius* membership using the *eff_size* function in the R package *emmeans* v 1.7.0. To test whether the scaling relationship between the visual and olfactory regions of the calyx differs between species, we used a GLM with olfactory calyx volume and species as fixed effects. The building of models and testing for pairwise differences follows the methods set out under section 2.2. We also assessed whether the scaling relationship between the visual and olfactory calyces differs between species using the function *sma* from the R package *smatr* v 3.4-8, testing for pairwise differences in both slope and elevation. The “robust” option was set to true for these analyses to minimise the influence of potential outliers (28).

Finally, the R package *bayou* v 2.0, was used to identify evidence of phylogenetic shifts in the scaling relationship between the visual and olfactory regions of the calyx within the eight sampled species (see 2.1.ii) (29). We compared three models: one with no shifts in either slope or elevation (all species are assumed exhibit the same allometric scaling relationship), one allowing for shifts in elevation (a common allometric slope is assumed, but “grade-shifts” are allowed), and one allowing for shifts in both elevation and slope. Models were each run for 1,000,000 iterations, with a burn-in of 300,000 generations and their fits were compared using the *steppingstone* function in *bayou*, run for 10,000 generations with a burn-in of 3000 (30). The posterior probability cut-off for identifying a shift in the relationship between mushroom body and rCBR size was set at 0.5 following published recommendations (29,31,32).

#### 2.4 Testing ecological hypotheses

To determine whether key shifts in mushroom body size co-occurred with transitions in pollen feeding state we estimated ancestral states (presence/absence of pollen feeding) in internal nodes. The state of the discrete trait of pollen feeding at internal nodes was estimated using three different methods: (1) MCMC stochastic character mapping in *phytools* for 1000 simulations, (2) maximum likelihood using the *ace* function in the R package *ape* v 5.5 (48), and (3) maximum parsimony using the *asr_max_parsimony* function in the R package *castor* v 1.7.0 (49). The *fitDiscrete* function from the R package *geiger* 2.0.7 (50) was used to determine the best fitting transition model between equal rates, symmetric and all rates differ. ‘Equal Rates’ was the best fitting model and was used for all downstream analyses.

Finally, we explored whether certain ecological factors could explain variation in mushroom body size across Heliconiini using phylogenetic generalised linear mixed models (GLMM) in *MCMCglmm* v 2.32 (24). We took the best fitting model explaining variation in mushroom body size (see 2.1) and then included ecological factors – the degree of social roosting, host plant number/generalism, and pollen feeding (which was tested by removing the *Heliconius* factor with which it is almost perfectly confounded), to test whether these improve the fit of the model. All models were checked for convergence using the *gelman.diag* function and for auto-correlation using the *autocorr* function provided in *MCMCglmm*, in addition to visually inspecting the trace plots. All models were run for 500,000 iterations, with a burn-in of 10,000 and a thinning factor of 500.

### 3. Behavioural experiments

#### 3.1 Animals

The behavioural experiments, using colour cues, were carried out on captive-reared butterflies between in either Gamboa, Panama, or, during travel restrictions related to the Covid-19 pandemic, in greenhouses at the University of Bristol, UK. All experiments were run using *Heliconius erato* and *Dryas iulia*, with additional *Heliconius melpomene* for the positive patterning experiment. All experiments were conducted using freshly-eclosed, naiive individuals.

In Gamboa, all larvae were reared from stocks established with locally caught, wild butterflies using the insectaries at the Smithsonian Tropical Research Institute in Gamboa, Panama. Stock butterflies were kept in 2 x 2 x 3 m mesh cages in ambient conditions with natural light. Larvae were reared in mesh pop-ups and were provided with fresh leaves daily. *H. erato* and *D. iulia* were reared on *P. biflora*, and *H. melpomene* on *P. triloba*. Training and testing of butterflies were conducted in 2 x 2 x 3 m mesh cages in ambient conditions under natural light. A single *Psychotria elata*, with all flowers removed, was placed in the rear right corner of these cages as a roosting site.

In Bristol, *D. iulia* and *H. erato* pupae were shipped to Bristol from the Stratford Butterfly Farm (UK) and Costa Rica Entomological Supply (Costa Rica). Pupae were kept in a climate-controlled greenhouse at 30°C and 80% humidity until emergence. The biconditional discrimination assay was conducted using adults that emerged from these pupae.

#### 3.2 Positive patterning

The positive patterning experiment was conducted at the Smithsonian Tropical Research Institute insectaries in Gamboa, Panama, 2019. For this assay, stock populations of *Dryas iulia*, *H. erato* and *H. melpomene* were established from wild-caught individuals around Gamboa, Panama. Stock populations were kept in outdoor cages in ambient conditions. Adult stock butterflies were fed daily with a sugar-protein solution (20% sugar, 5% Vertark Critical Care Formula, 75% water, w/v) mixture and had access to *Psiguria* and *Lantana* flowers. Larvae of both species were fed with *Passiflora biflora* leaves. The positive patterning assay was conducted using only captive-reared butterflies from these stock populations. The day after eclosion, experimental individuals were introduced to a pre-training cage containing only white feeders filled with sugar-protein solution. Individuals were kept in this pre-training environment for two full days to acclimatise to using the feeders before beginning training.

After pre-training, butterflies were subject to an initial preference test between three types of artificial feeders: yellow, purple, yellow + purple. Butterflies were introduced to a testing cage containing 12 artificial feeders (4 yellow, 4 purple and 4 yellow + purple) arranged randomly with at least 6.5 cm between feeders on each side. To ensure that butterflies responded to visual cues only, feeders in the testing cages were empty. Butterflies were deprived of food from 12:00 the day prior to testing to encourage feeding during the trials. The preference test lasted for four hours and was filmed using mounted GoPro Hero 5 cameras. The film was then reviewed to count the number of feeding attempts per individual on each colour. A feeding attempt was only counted if the butterfly landed on the feeder and probed it with its proboscis.

After the initial preference test, butterflies were placed in training cages containing 12 feeders (4 yellow, 4 purple and 4 yellow + purple). The yellow + purple feeders contained a food reward while the yellow and purple feeders were filled with a saturated solution of quinine, serving as an aversive stimulus. These feeders were arranged randomly each morning. Butterflies kept in training cages for eight days and could freely sample the feeders for the entire period. Following the training period, butterfly feeding preferences were re-tested, following the same protocol as the initial preference test.

#### 3.3. Biconditional discrimination

The biconditional discrimination trials were conducted in greenhouses at the University of Bristol in 2020. For this assay. The biconditional discrimination assay was conducted using adults that emerged from these commercially sourced pupae. The day after eclosion experimental individuals were introduced to a pre-training cage containing only white feeders filled with sugar-protein solution. Individuals were kept in this pre-training environment for two full days to acclimatise to using the feeders before beginning training.

The biconditional discrimination assay used artificial feeders of four different colour combinations: red + blue, purple + yellow, red + yellow, purple + blue (Figure S1). Following the pre-training period, butterflies were placed in a cage containing four empty feeders of each colour combination to test for their initial preference, following the testing protocol described for the positive patterning assay. Butterflies were then randomly assigned to one of two training regimes.

For the first training regime, butterflies were introduced to a cage containing four feeders of each colour combination, with the purple + yellow and blue + red feeders containing a food reward and the yellow + red and blue + purple combination filled with the aversive quinine solution (Figure S1). Each colour was therefore evenly represented between rewarded and punished feeders. Accordingly, butterflies could not solve the task by learning a single colour but needed to learn specific combinations of colours. The training regime for the second group was the reverse of the first, with yellow + red and purple + blue feeders rewarded and purple + yellow and blue + red punished. The eight-day training period and subsequent testing followed the same protocol as the positive patterning assay.

**Figure S1:**
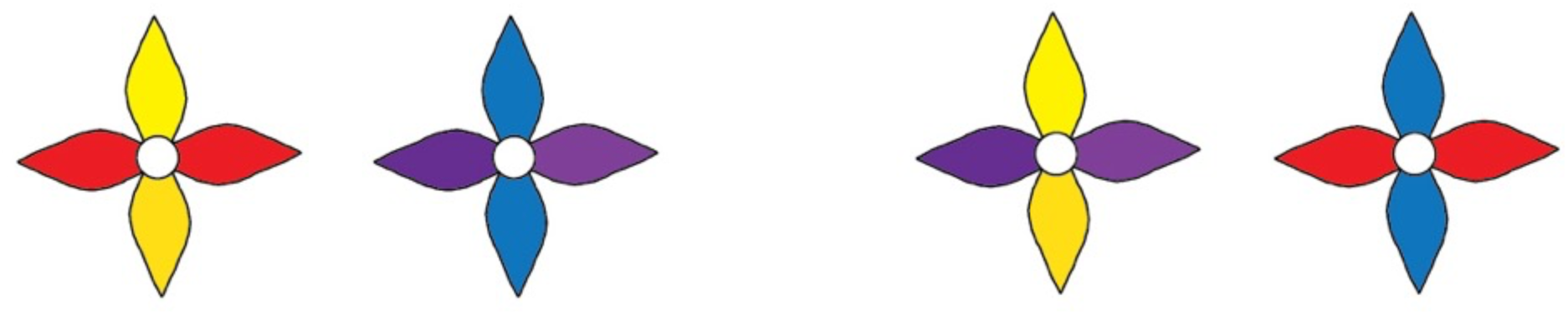
The four colour combinations used during the biconditional discrimination trial. These pairs of combinations were presented with either a food reward (sugar-protein) or aversive stimulus (quinine). Each colour was represented in two different combinations.

#### 3.4 Long term memory

Long-term memory (LTM) and reversal learning (RL) experiments, using colour cues, were carried out on captive-reared butterflies between January and April 2019 in Gamboa, Panama. Individuals were transferred to a pre-training cage one day after eclosion. Here, butterflies were fed solely with white artificial feeders containing a sugar-protein solution (20% sugar, 5% Vertark Critical Care Formula, 75% water, w/v) for two days (from 08:00 to 12:00) to familiarise them with the use of artificial feeders.

After pre-training, butterflies were introduced to a testing cage to determine initial feeding preferences between purple and yellow. Testing cages contained 12 purple and 12 yellow feeders randomly in a 4 X 6 grid, with 6.5 cm between feeders on each side. To ensure that butterflies responded exclusively to visual cues, feeders in the testing cages were empty. Preference testing lasted for four hours from 08:00 to 12:00 and was filmed from above using a GoPro Hero 5 camera mounted on a tripod. Butterflies were individually numbered on their wings for identification using a permanent marker. The film was then reviewed to count the number of feeding attempts per individual on each colour, with up to 30 attempts recorded per individual. A feeding attempt was only counted if the butterfly landed on the feeder and probed it with its proboscis.

Butterflies were then trained to associate a food reward with their non-favoured colour, based on the results of their initial preference test. For butterflies that initially preferred purple, the training cage contained yellow feeders containing a sugar-protein solution, and purple feeders containing a saturated quinine solution, an aversive stimulus. The opposite arrangement was employed for individuals that initially preferred yellow. This training period lasted for four full days. After training, butterfly preferences were re-tested, following the same protocol as the initial preference test, to verify that individuals had indeed acquired the colour-food association.

After the trained preference test, individuals participating in the long-term memory assay were placed for eight days in a cage identical to the pre-training cage, containing only white feeders filled with a sugar-protein solution. The deprivation of colour stimuli for eight days allowed for testing the long-term memory retention of the colour-food association acquired during the training period, and ensured that long-term memory was being tested rather than short-term or mid-term memory. A period of eight days was chosen because *Heliconius* are known to maintain their foraging routes over periods ranging from weeks to months, during which time a pollen resource could be unproductive for several days due to competition or damage, but ultimately rewarding over the long term. Butterflies were then subject to a third preference test to determine if the learned preference was maintained, following the same protocol as the initial preference test. When subsequently analysing the data, individuals that exhibited less than 50% accuracy during the initial trained preference test, and therefore did not appear to have learned the food-colour association, were removed from the dataset. In total, 1 out of 48 *H. erato* individuals, and 4 of 63 *Dryas iulia,* were removed from the dataset.

#### 3.5 Statistical analysis

Learning performance in the positive patterning and biconditional discrimination assays was analysed with generalised linear mixed models (GLMMs) using the *glmer* function from the R package *lme4* v 1.1-27.1 (51). These models included species and training as fixed effects (in addition to their interaction), with an individual-level random effect. For the biconditional discrimination assay, since different butterflies were trained to different colour combinations, training regime was also included as a fixed effect. However, training regime was non-significant and therefore removed from the model. To test for interspecific differences in the drop in performance between the initial preference test and the long-term memory test, an interaction between species and trial was also included. Diagnostics for these models were assessed using the package *DHARMa* (46). All post hoc comparisons were made by obtaining the estimated marginal means using the R package *emmeans* v 1.7.0 and were corrected for selected multiple comparisons using the Tukey test (47).

### 4. Supplementary Results

#### 4.1 Sex effects on mushroom body size

*Heliconius* females have significantly larger mushroom bodies than males (pMCMC < 0.001), whereas no such sex effect was identified in the outgroup Heliconiini (pMCMC = 0.690) (Figure S2). This difference could potentially reflects sex differences in pollen foraging behaviour, with females tending to collect more pollen than males (52, 53).

**Figure S2:**
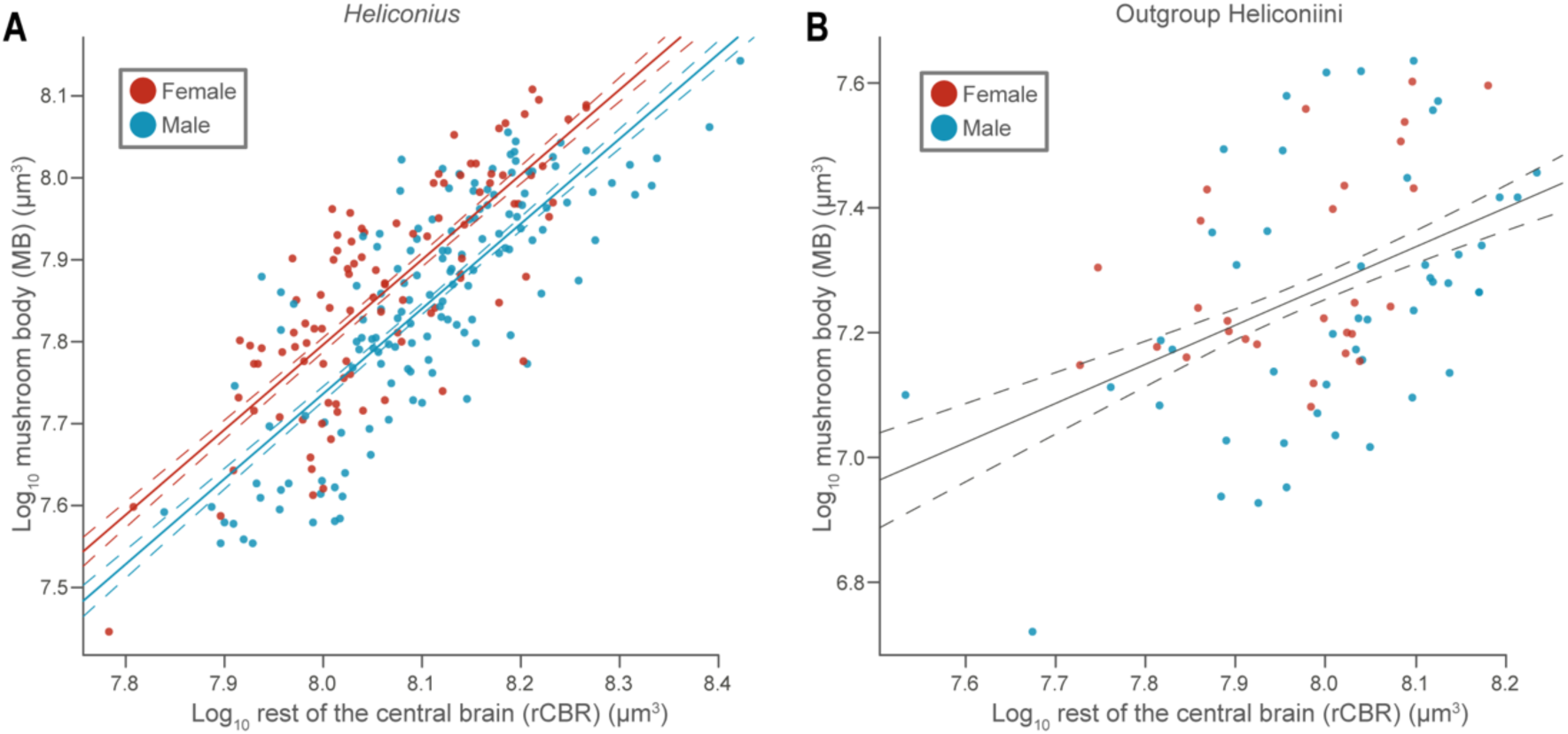
Sexual dimorphism in mushroom body size in *Heliconius*. (**A**) Female *Heliconius* have larger mushroom bodies than males (pMCMC < 0.001). (**B**) There is no sex effect for outgroup Heliconiini (pMCMC = 0.690). Data points represent individuals (n_Heliconius_ = 243; n_Outgroup Heliconiini_ = 75). Dashed lines show standard error.

#### 4.2 Clade and pairwise differences in mushroom body scaling

Within the Heliconiini, mushroom body expansion did not occur solely along the branch leading to *Heliconius*, but rather occurred in a stepwise series along the phylogeny (Figure S3). Among the non-*Eueides* and non-*Heliconius* Heliconiini, *Dryadula phaetusa* shows an expansion of the mushroom bodies (Figure S3A). The entire *Eueides* genus shows mushroom body expansion comparable to that in *Dryadula phaetusa*, with a further expansion in the ‘*lybia’* subclade (Figure S2B). This culminates in a further, even more marked, expansion in *Heliconius* (Figure S3C). Notably, the loss of pollen feeding in *Heliconius aeode* is not associated with a corresponding decrease in mushroom body volume (Figure S3C; yellow). GLMM analysis that include phylogenetic grouping (outgroup genus or subclade within *Heliconius*) as a factor shows mushroom body size, controlling for rCBR, antennal lobe and medulla size, as internally consistent both within *Heliconius* and amongst the outgroup Heliconiini (Table S8). However, MB volume in *Dryadula phaetusa* is not significantly different to either the *Heliconius* clades or the other outgroups, while *Eueides* only differ from *melpomene*-group *Heliconius* (Table S8). Uncorrected pairwise comparisons, however, do show *Dryadula phaetusa* and *Eueides* as having significantly larger MBs than several outgroup Heliconiini, and smaller MBs than some Heliconius.

Supporting these results, pairwise allometric analysis with the R package *smatr* indicates that while the slope of the allometric relationship between rCBR and mushroom body volume does not differ between Heliconiini species, there are significant differences in elevation (Table S9). In general, relative mushroom body volume is consistent throughout *Heliconius* and larger than in the outgroups, congruent with the results of the phylogenetic GLMM above. *Eueides* and *Dryadula phaetusa* have MB volumes intermediate between *Heliconius* and the other outgroups, while *H. telesiphe*, *H. doris* and *H. aoede* have smaller mushroom bodies than several other *Heliconius* species.

**Figure S3:**
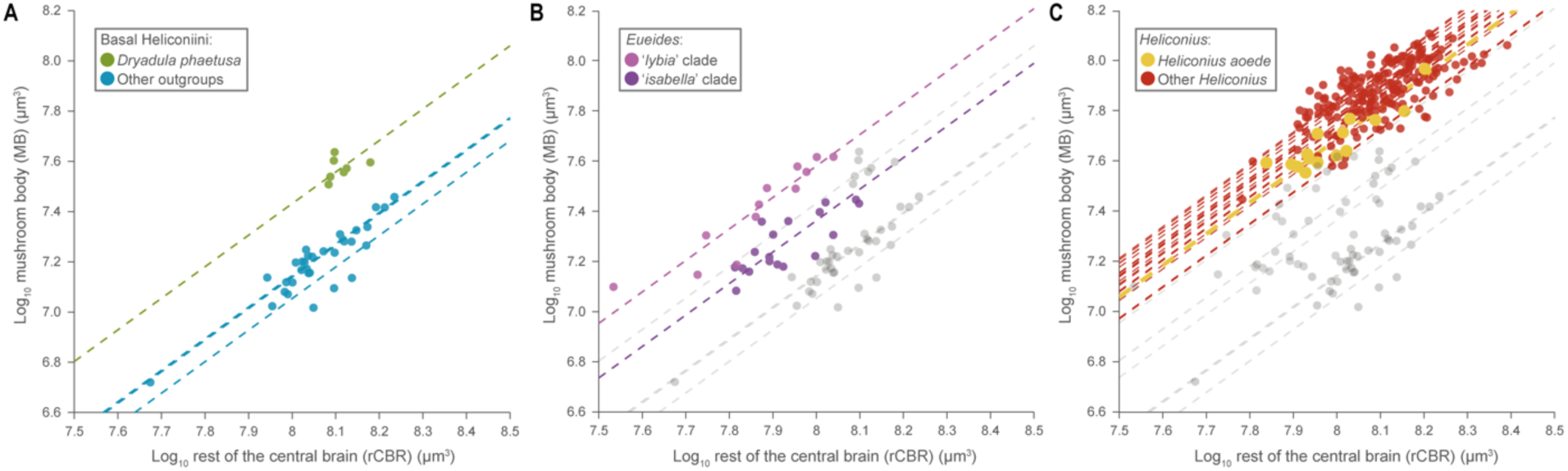
Stepwise increases in mushroom body (MB) across the Heliconiini phylogeny. (**A**) Amongst the non-*Heliconius* and non-*Euedies* Heliconiini, *Dryadula phaetusa* show enlarged MBs. (**B**) *Eueides* show an upshift in MB size relative to non-*Dyradula* basal Heliconiini and within the genus, the ‘*lybia’* clade shows a further enlargement. (**C**) *Heliconius* show a further, even larger, increase in mushroom body size. Notably, the non-pollen-feeding *Heliconius aoede* (yellow) does not show an decrease in mushroom body size relative to other *Heliconius*. Regression lines show (A,B) species and (B) main clades. Points indicate individuals.

**Table S8.**
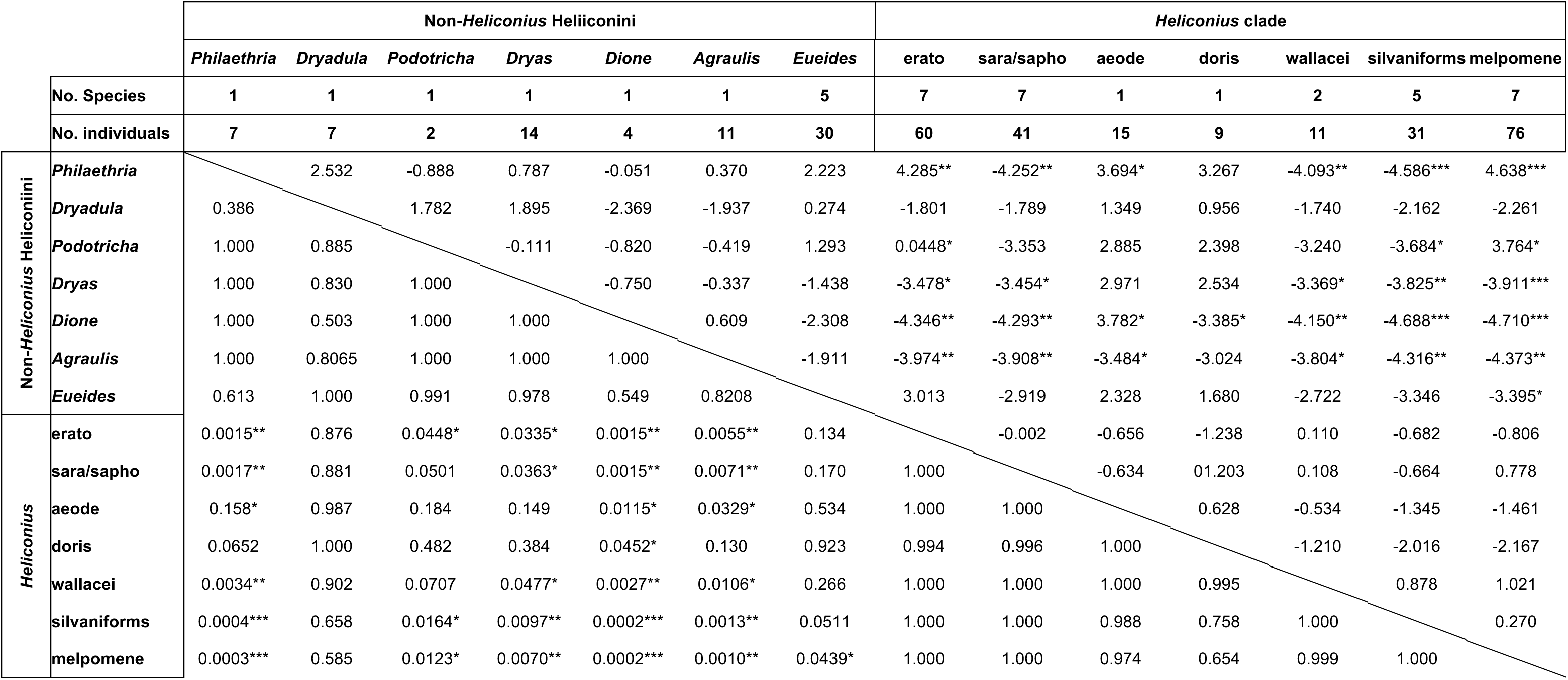
Pairwise comparisons between Heliconiini clades for mushroom body size, derived from a phylogenetic generalised linear mixed model including the size of the antennal lobe, the medulla and the rest of the central brain as fixed effects. Z-ratio shown above the diagonal and p-values below, corrected for multiple comparisons using Tukey’s test (* = p<0.05; ** = p<0.01; *** = p<0.001).

**Table S9.**
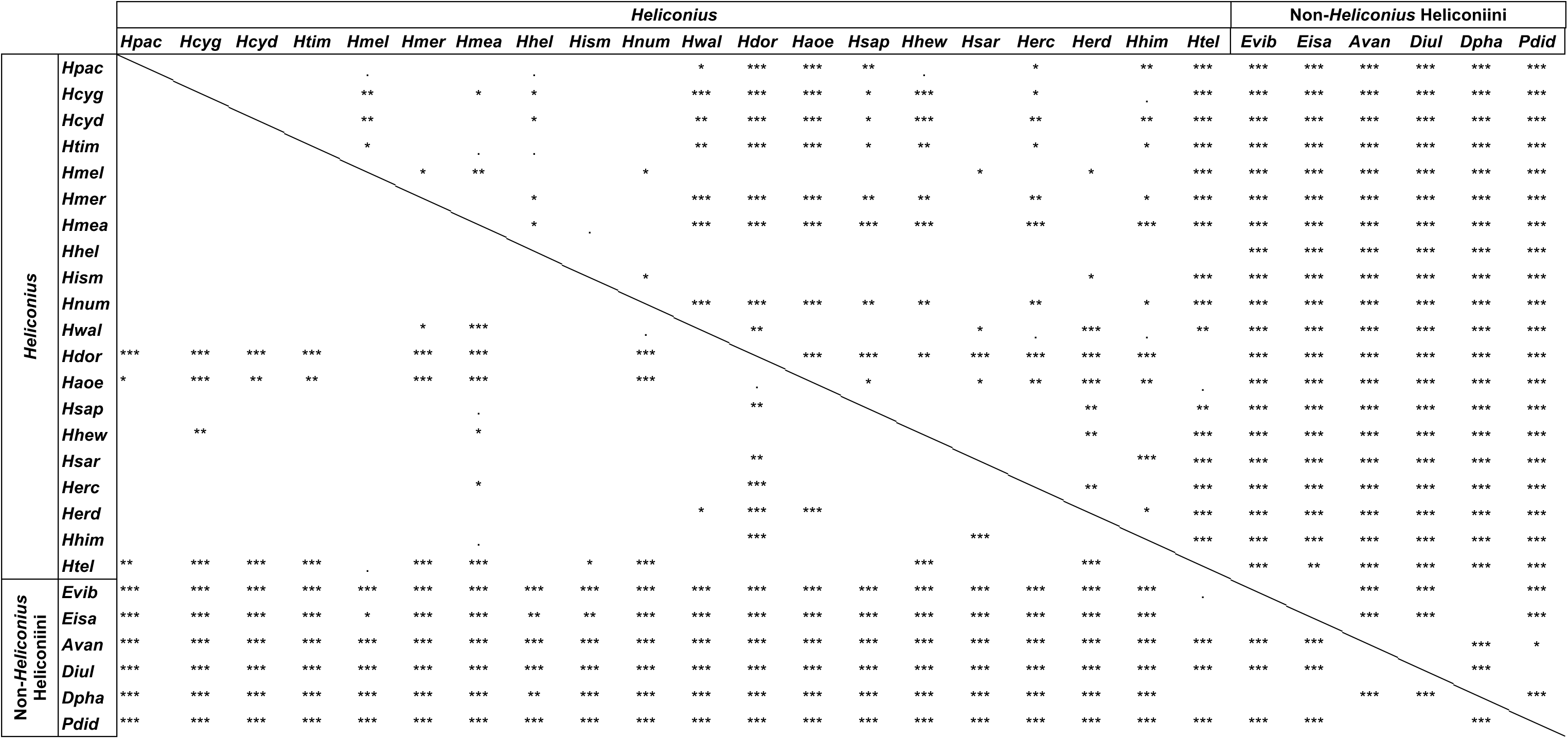
Interspecific differences in elevation in the scaling relationship between the size of the mushroom bodies and the rest of the central brain. P-values above the diagonal are uncorrected, and below are corrected for multiple comparisons. (. = p < 0.01; * = p < 0.05; ** = p < 0.01; *** p < 0.001; *Avan* = *Agraulis vanillae*; *Diul* = *Dryas iulia; Dpha* = *Dryadula phaetusa*; *Eisa* = *Eueides Isabella*; *Evib* = *Eueides vibilia*; *Haoe* = *H. aoede*; *Hcyd* = *H. cyndo chioneus*; *Hcyg* = *H. cydno galanthus*; *Hdor* = *H. doris*; *Herc* = *H. erato cyrbia*; *Herd* = *H. erato demophoon*; *Hhel* = *H. hecale*; *Hhew* = *H. hewitsoni*; *Hhim* = *H. himera*; *Hism* = *H. ismenius*; *Hmea* = *H. melpomene amaryllis*; *Hmel* = *H. melpomene melpomene*; *Hmer*; *H. melpomene rosina*; *Hnum* = *H. numata*; *Hpac* = *H. pachinus*; *Hsap* = *H. sapho*; *Hsar* = *H. sara*; *Htel* = *H. telesiphe*; *Htim* = *H. timareta*; *Hwal* = *H. wallacei*; *Pdid* = *Philaethria dido*).

#### 4.3 Additional results on mushroom body evolutionary rates

Our findings of a marked expansion the mushroom body in *Heliconius* are corroborated by our re-analysis of an independently collected neuroanatomical dataset of 41 species of North American butterflies, including *H. charithonia* and the Heliconiini *Agraulis vanillae* (54). Using the R package *bayou v* 2.0, we identified phylogenetic shifts in the scaling relationship between the mushroom body and the rest of the central brain. The best fitting model permitted shifts in elevation specifically (marginal likelihoods: elevation shifts = 25.874; both = 7.003; none = 7.656) and identified two shifts with a posterior probability greater than 0.5 (Figure 2A,B). We resolved a distinct upshift in mushroom body volume in *Heliconius* (post. prob. = 0.90; Figure S4; Table S11), despite an overall decrease in mushroom body volume in the Nymphalidae as a whole (excluding *Danaus plexippus*) (post. prob. = 0.69; Figure S4A,B; Table S11). Similarly, *H. charithonia* stands out amongst the sampled butterflies as having an exceptionally elevated rate of mushroom body evolution when analysed using both the *multirateBM* function in the R package *phytools* (Figure S4C,D) and the variable rates model in *BayesTraits* v 3 (Bayes Factor = 3.542; Figure S5C,D).

**Figure S4:**
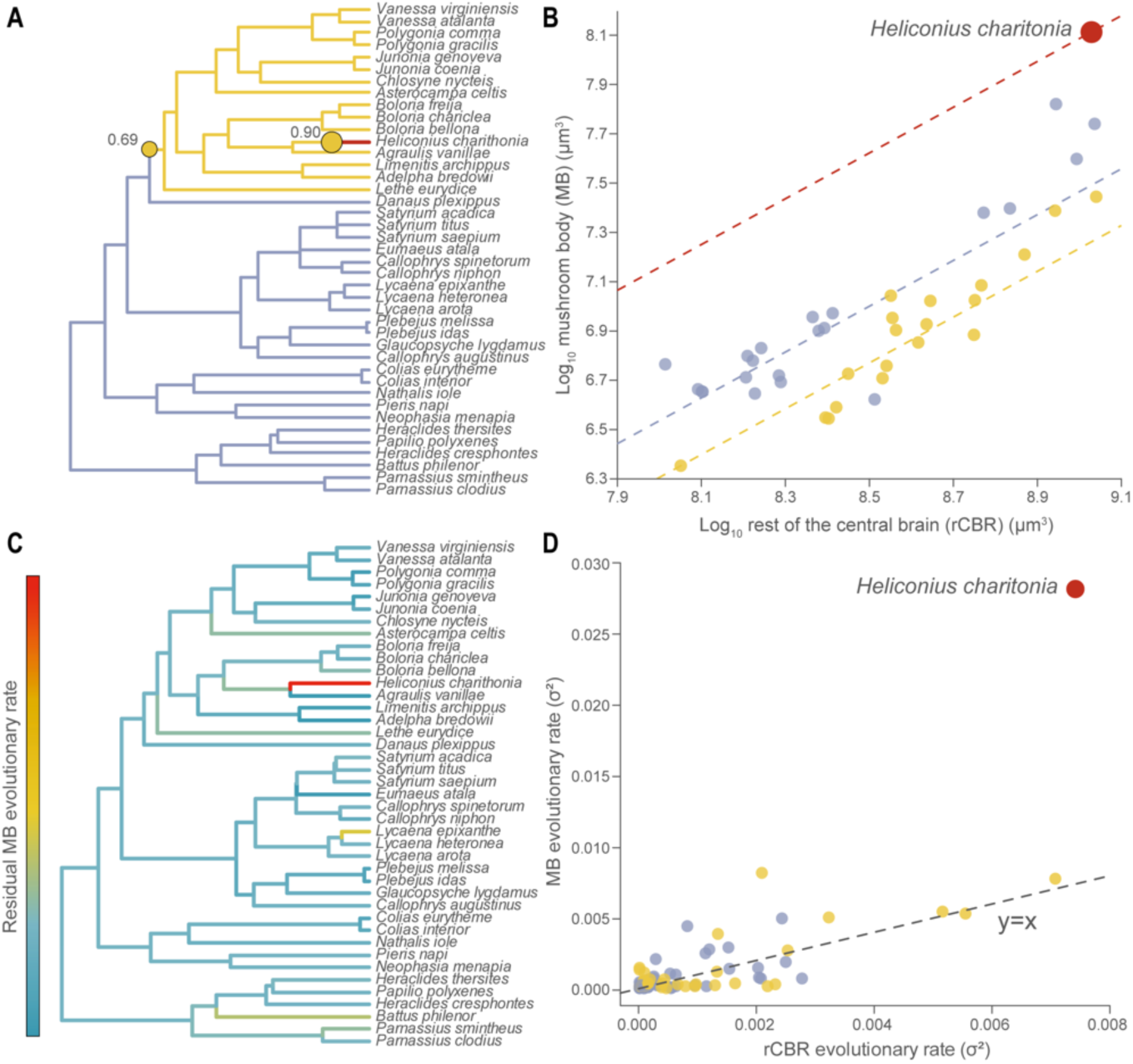
*Heliconius* stand out as having markedly expanded and increased evolutionary rate of the mushroom bodies even in a phylogenetically wide sampling of butterflies. (**A** and **B**) Phylogenetic shifts in the scaling relationship between the volume of the mushroom body (MB) and the rest of the central brain (rCBR) across 41 North American butterfly species (posterior probability > 0.5). Despite a decrease in MB volume in the Nymphalidae (excluding *Danaus plexippus*) (blue), *Heliconius* exhibit a dramatic increase in MB volume relative to this phylogenetically wide sampling of butterflies. (**C** and **D**) This increase in MB size is associated with a pronounced increase in the estimated evolutionary rate of MB volume, controlling for the evolutionary rate of rCBR volume.

**Figure S5:**
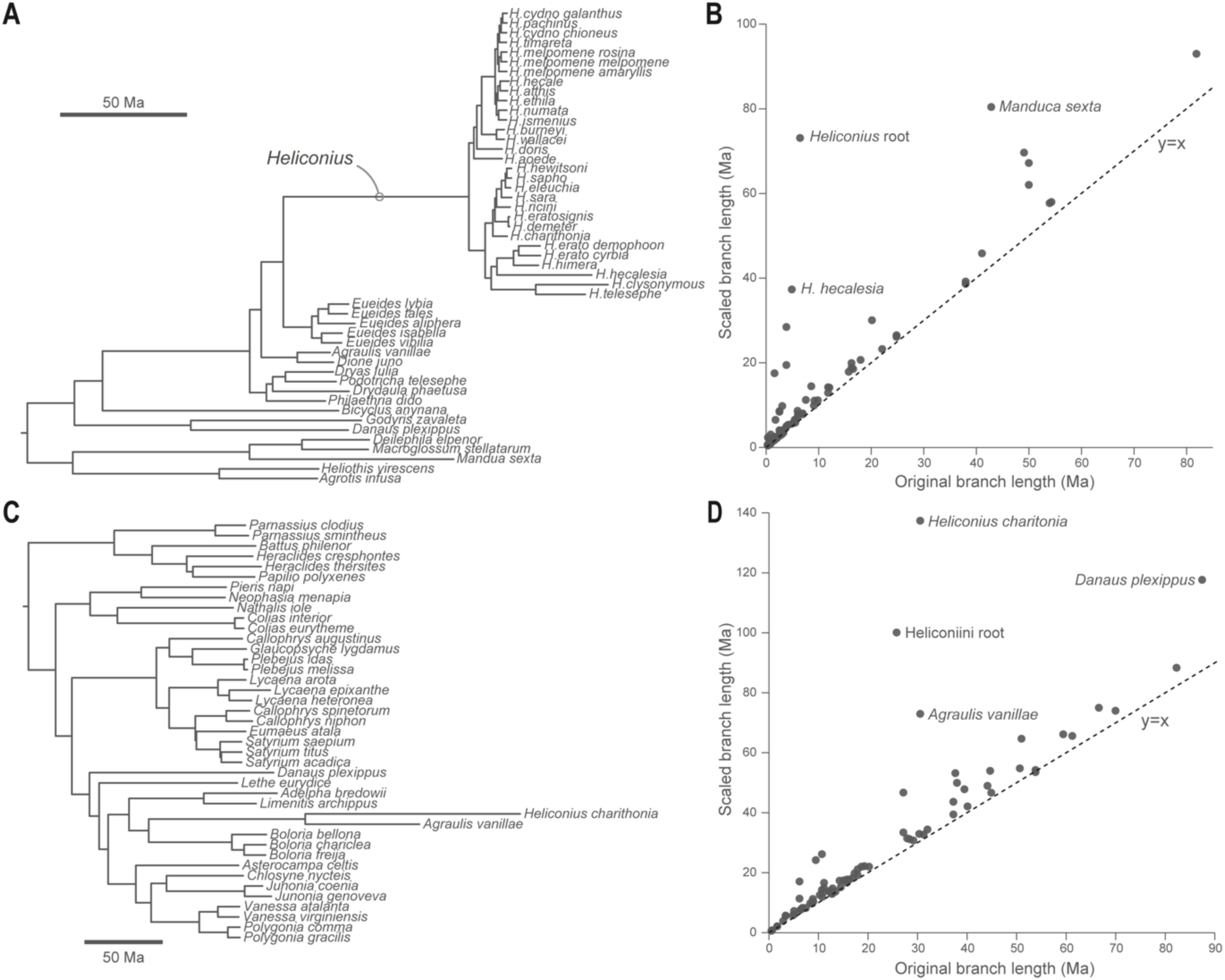
Increased evolutionary rate of the mushroom body in *Heliconius*. Estimates of the evolutionary rate of the mushroom body (MB) (controlling for the size of the rest of the central brain) produced from multiple iterations of a variable rates models in *BayesTraits* v 3, using originally ultrametric trees. (**A**) Consensus scaled tree for 41 Heliconiini species and eight outgroup Lepidoptera, with (**B**) scaled branch length plotted against original branch lengths. (**C**) Consensus scaled tree for 41 North American butterfly species, including *H. charithonia* and *Agraulis vanilla*, generated using neuroanatomical data from Snell-Rodd et al. (2020) and the consensus tree in Earl et al. (2021), with (**D**) scaled branch lengths plotted against original branch lengths. Increased branch lengths in (A) and (C) and deviance from y = x in (B) and (D) indicate an estimated increase in the evolutionary rate of the MB.

#### 4.4 Additional neuroanatomical detail

**Figure S6:**
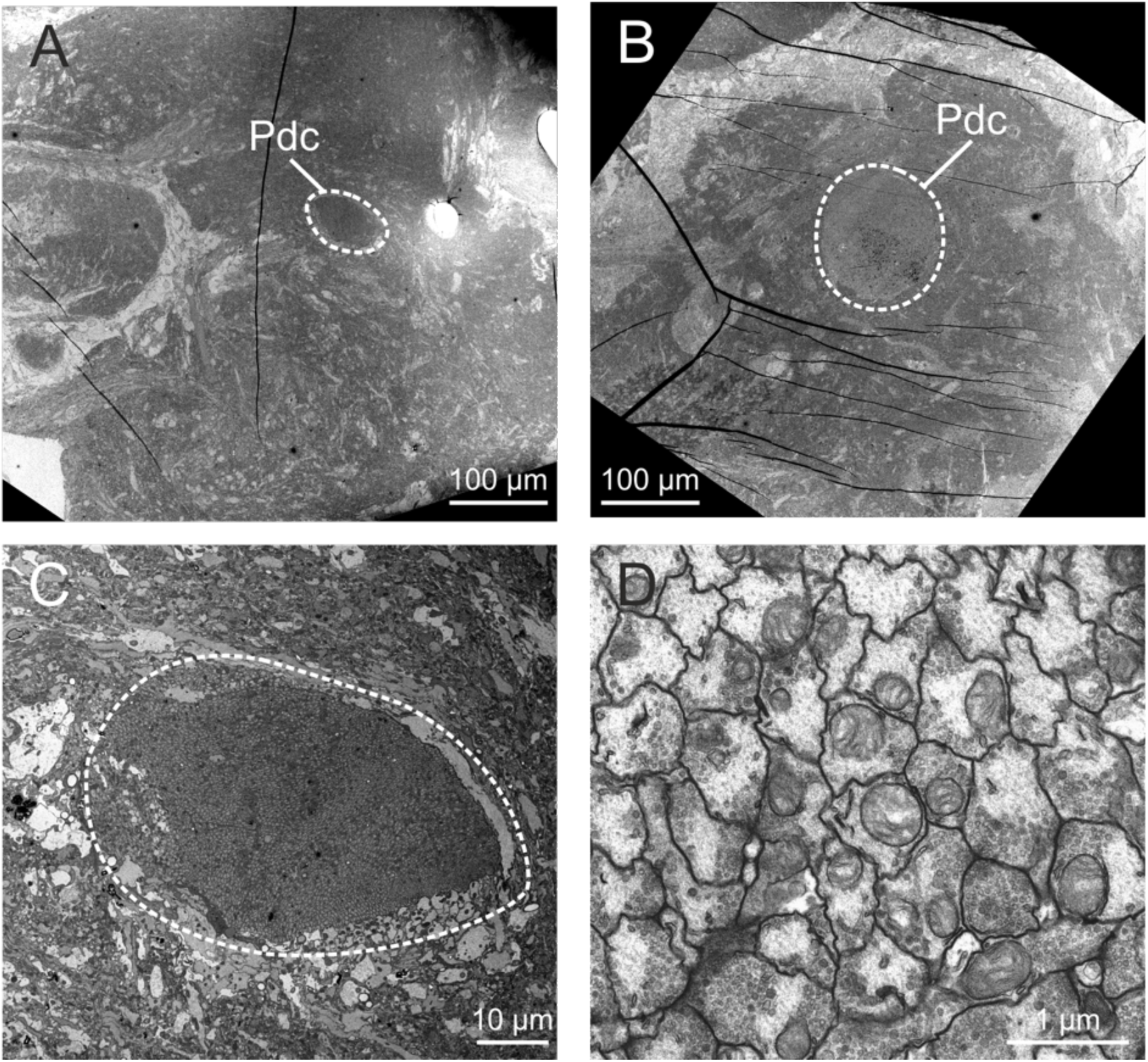
An example of the electron microscopy imaging. **(A)** low magnification of orthogonally sectioned brains showing the peduncle (Pdc) in *Dryas iulia*, and **(B)** *Heliconius erato*. **(C)** Mid-magnification cross section through at *Dryas iulia* and **(D)** high magnification image of cross-sectioned Kenyon cell axons running through the peduncle. Total Kenyon cell counts were estimated by measuring the area of the cross section at mid magnification and the density of Kenyon cell axons at high magnification, and multiplying the two estimates.

**Figure S7:**
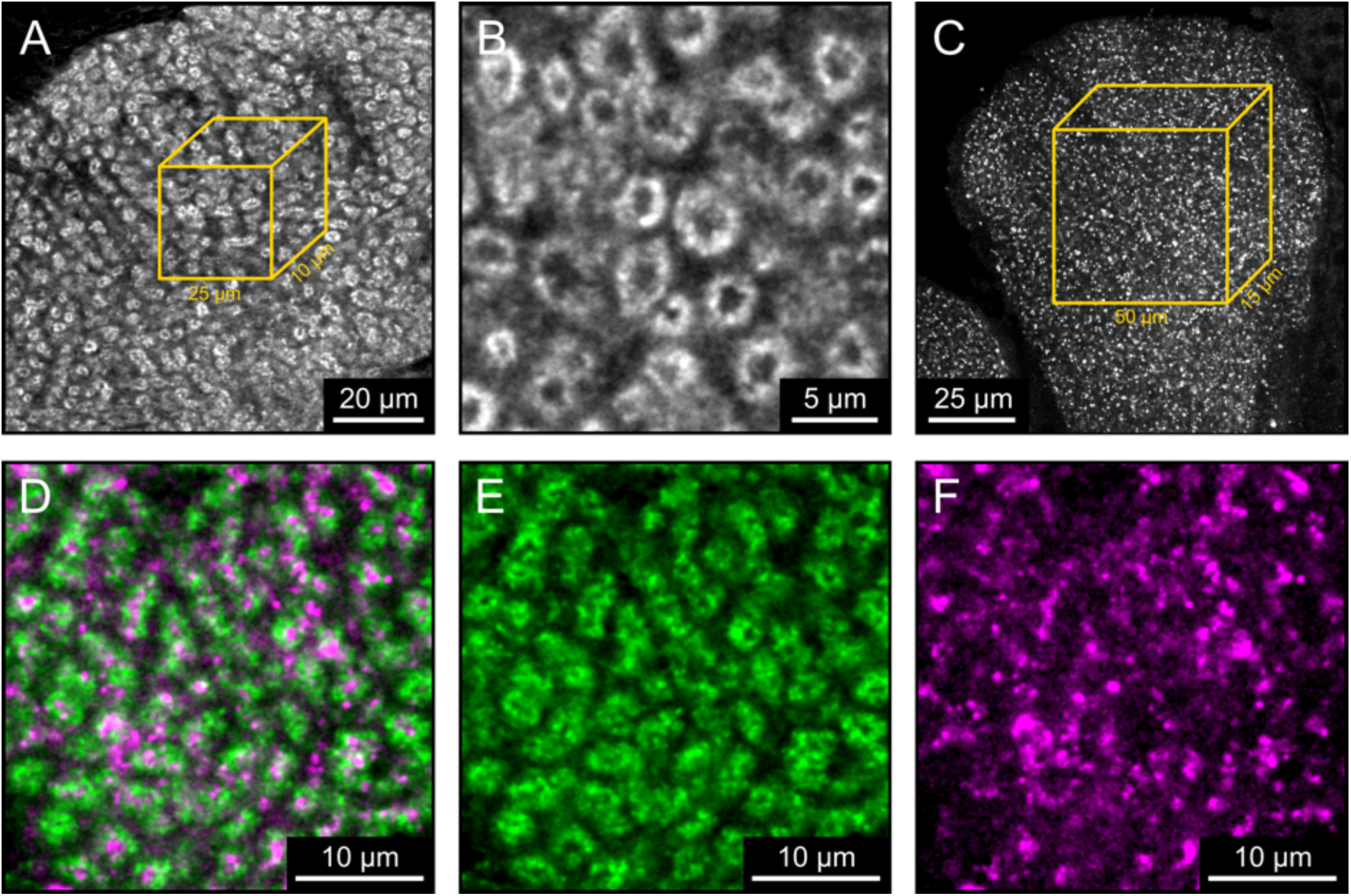
An example of the microglomeruli staining. **(A)** An optical section through a portion of the calyx of *H. charithonia* after staining of the Kenyon cells dendrites with phalloidin conjugated dye. A cube of 25×25×10 µm was extracted from the image stack to manually count the microglomeruli structure at **(B)** high magnification. **(C)** An optical section through a portion of the calyx of *H. hecale* after staining of the synaptic boutons with anti-Synorf1 antibody. Synaptic boutons were automatically counted within cubes of 50×50×15 µm. **(D)** Combined labelling of **(E)** Kenyon cell dendrites and **(F)** synaptic terminals in the calyx of *H. erato*. The microglomeruli structure appear to be innervated by more than a single postsynaptic bouton.

**Figure S8:**
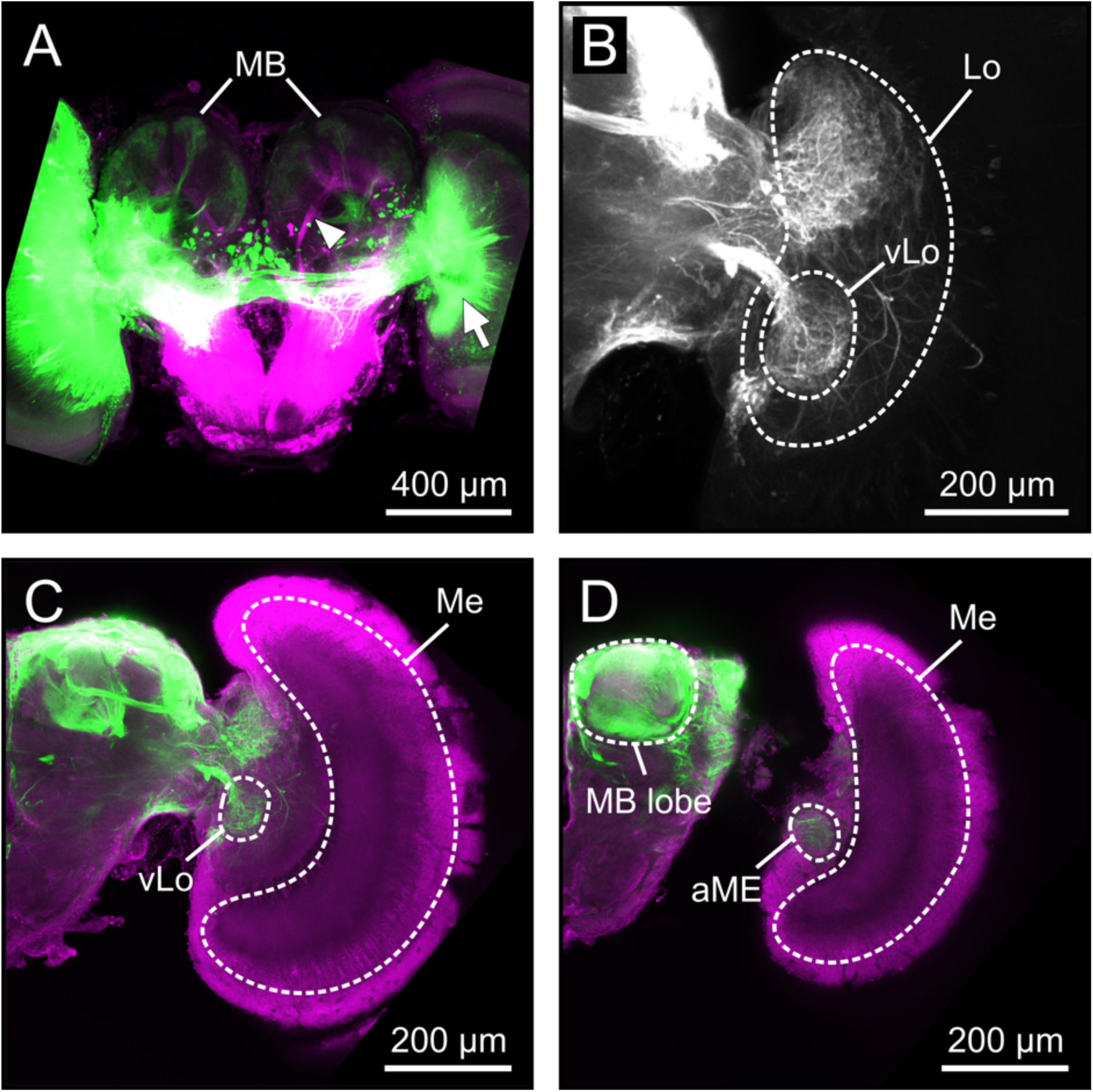
Identifying visual neuropils that innervate the mushroom body calyx. **(A)** Optical section through the brain of *H. melpomene* after injection of a fluorescent tracer in the optic lobes (green) and the antennal lobe (magenta). Both sensory neuropil have projection into the mushroom bodies. The arrowhead shows a major track of olfactory projection neurons, whereas the complete arrow shows the injection site in the optic lobes. **(B)** An optical section through the optic lobe after injection of a fluorescent tracer in the mushroom body calyx itself. The visual calyx is mainly arborized by projection neurons from the ventral lobe of the lobula (vLo). **(C)** A large tract of visual projection neurons leave the vLo as ab optical relay and run through the central brain to the MB. **(D)** At the dorsal surface, the accessory medulla (aMe) also send projection to the MB calyx.

**Table S10.** Interspecific differences in total Kenyon cell number. Pairwise comparisons from generalised linear model testing the effect of species on Kenyon cell number, corrected for multiple comparisons using Tukey’s test. T-ratio values shown above the diagonal and p-values below. (* = p <0.05; ** = p < 0.01; *** = p < 0.001).

|  | *Dryas iulia* | *Agraulis vanillae* | *E. isabella* | *H. hecale* | *H. melpomene* | *H. hortense* |
| --- | --- | --- | --- | --- | --- | --- |
| *Dryas iulia* |  | -2.159 | -7.833*** | -21.324*** | -22.868*** | -23.201*** |
| *Agraulis vanillae* | 0.2774 |  | -10.416*** | -24.636*** | -26.263*** | -26.614*** |
| *E. isabella* | <0.0001*** | <0.0001*** |  | -13.491*** | -15.035*** | -15.368*** |
| *H. hecale* | <0.0001*** | <0.0001*** | <0.0001*** |  | -1.544 | -1.877 |
| *H. melpomene* | <0.0001*** | <0.0001*** | <0.0001*** | 0.6385 |  | 0.333 |
| *H. hortense* | <0.0001*** | <0.0001*** | <0.0001*** | 0.4294 | 0.9994 |  |

**Table S11.** Interspecific differences in the scaling relationship between Kenyon cell number and calyx volume. Pairwise comparisons from generalised linear model testing the effect of species on mushroom body calyx volume, controlling for Kenyon cell number, corrected for multiple comparisons using Tukey’s test. T-ratio values shown above the diagonal and p-values below. (* = p <0.05; ** = p < 0.01; *** = p < 0.001).

|  | *Dryas iulia* | *Agraulis vanillae* | *E. isabella* | *H. hecale* | *H. melpomene* | *H. hortense* |
| --- | --- | --- | --- | --- | --- | --- |
| *Dryas iulia* |  | 3.315* | 4.115* | -0.144 | 1.598 | 0.112 |
| *Agraulis vanillae* | 0.0215* |  | 1.732 | -1.039 | 0.627 | -0.739 |
| *E. isabella* | 0.0022* | 0.5187 |  | -3.023* | -0.255 | -2.361 |
| *H. hecale* | 1.0000 | 0.9020 | 0.0453* |  | 6.058*** | -0.859 |
| *H. melpomene* | 0.6044 | 0.9884 | 0.9998 | <0.0001*** |  | 5.320*** |
| *H. hortense* | 1.0000 | 0.9758 | 0.1929 | 0.9541 | <0.0001*** |  |

**Table S12.**
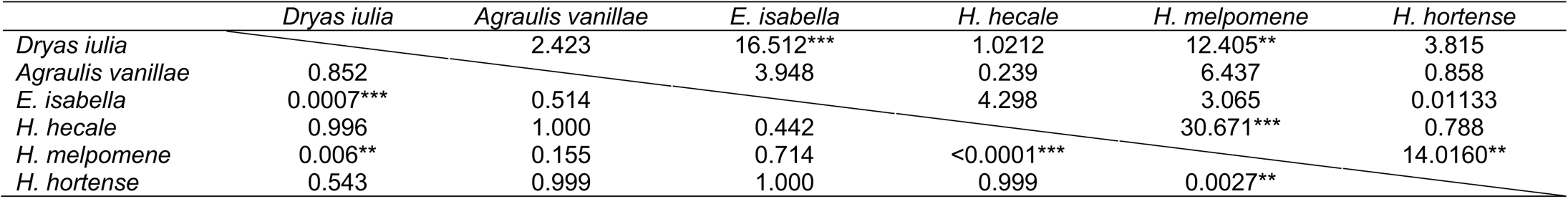
Interspecific differences in the scaling relationship between Kenyon cell number and calyx volume using SMATR. Pairwise comparisons for SMATR analysis testing for differences in elevation in the relationship between KC number and calyx volume, corrected for multiple comparisons. Test stat values shown above the diagonal and p-values below. (* = p <0.05; ** = p < 0.01; *** = p < 0.001).

#### 4.5 Variation in sensory innervation to the mushroom body calyx

**Table S13.** Pairwise comparisons from generalised linear model testing the effect of Species on visual calyx volume, controlling for olfactory calyx volume, corrected for multiple comparisons using Tukey’s test. T-ratio values shown above the diagonal and p-values below. * indicates p<0.05, ** indicates p<0.01, *** indicates p<0.001.

|  | *Dryadula phaetusa* | *Dryas iulia* | *Agraulis vanillae* | *E. isabella* | *H. doris* | *H. hecale* | *H. melpomene* | *H. hortense* |
| --- | --- | --- | --- | --- | --- | --- | --- | --- |
| *Dryadula phaetusa* |  | 5.658*** | 7.490*** | 2.994 | -12.981*** | -16.843*** | -19.083*** | -19.345*** |
| *Dryas iulia* | <0.0001*** |  | 3.352* | -3.501* | -16.835*** | -16.423*** | -18.028*** | -17.767*** |
| *Agraulis vanillae* | <0.0001*** | 0.0297* |  | -5.951*** | -17.537*** | -16.505*** | -17.930*** | -17.630*** |
| *E. isabella* | 0.0741 | 0.0197* | <0.0001*** |  | -15.787*** | -17.550*** | -19.563*** | -19.448*** |
| *H. doris* | <0.0001*** | <0.0001*** | <0.0001*** | <0.0001*** |  | -4.501*** | -6.599*** | 7.596*** |
| *H. hecale* | <0.0001*** | <0.0001*** | <0.0001*** | <0.0001*** | 0.0009*** |  | -2.474 | 4.246** |
| *H. melpomene* | <0.0001*** | <0.0001*** | <0.0001*** | <0.0001*** | <0.0001*** | 0.2285 |  | 1.801 |
| *H. hortense* | <0.0001*** | <0.0001*** | <0.0001*** | <0.0001*** | <0.0001*** | 0.0021** | 0.6222 |  |

**Table S14.**
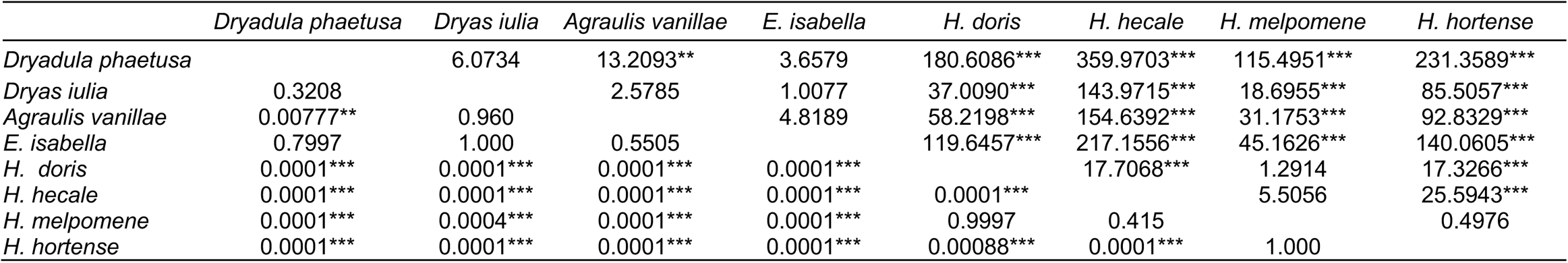
Pairwise comparisons for SMATR analysis testing for differences in elevation in the relationship between visual calyx volume and olfactory calyx volume, corrected for multiple comparisons. * indicates p<0.05, ** indicates p<0.01, *** indicates p<0.001.

#### 4.6 Evolution of ventral lobula size

Using the R package *bayou v* 2.0, we tested for phylogenetic shifts in the scaling relationship between the ventral lobula and the rest of the central brain. Although the best fitting model permitted shifts in elevation specifically (marginal likelihoods = 19.434) this was only marginally higher than the null model (marginal likelihood = 18.797) resulting in only weak evidence supporting these grade shifts (Bayes Factor = 1.274). Nevertheless, the model allowing for shifts in intercept did not identify any shifts associated with *Heliconius*, but rather size reductions of the ventral lobula at the base of *Eueides* and the *Philaethria dido* branch (Figure 2A,B). These size reductions appear to be associated with a slight increase in the evolutionary rate of the ventral lobula in *Eueides* (Figure S9C). *H. eratosignis* and *H. demeter* do show elevated rates of ventral lobula evolution (Figure S9C), but this is likely due to a minor, but significant, divergence in ventral lobula volume over very short branch lengths, as they otherwise cluster within *Heliconius* in an allometric plot (Figure 9B).

**Figure S9:**
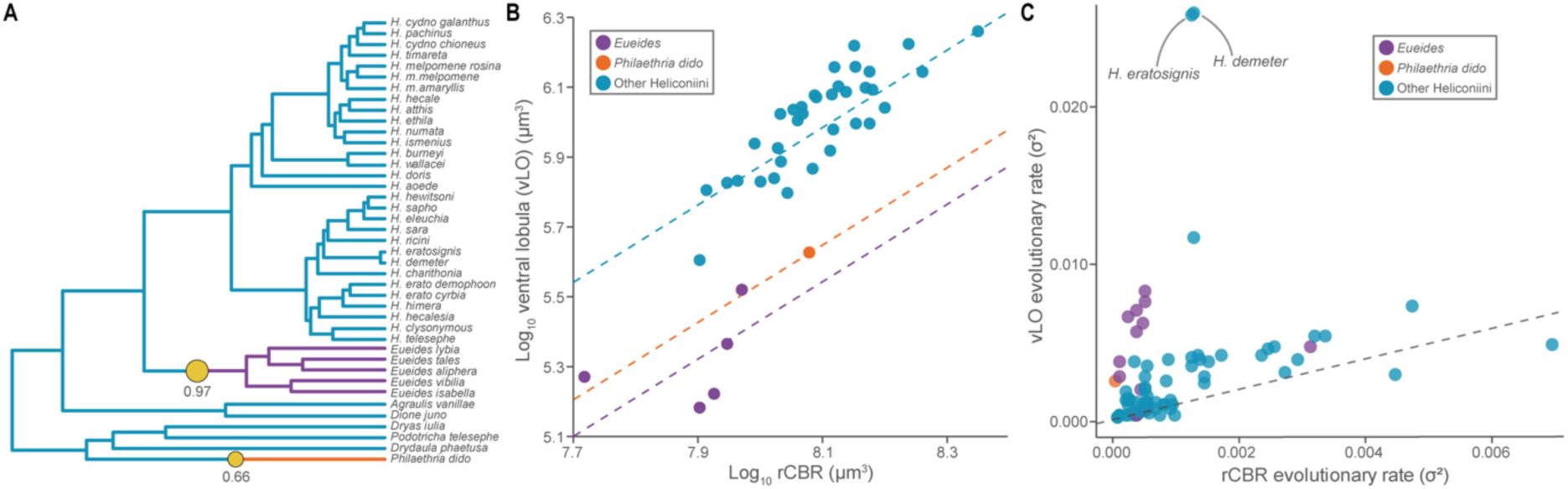
Variation in the volume and evolutionary rate of the ventral lobula across the Heliconiini. (**A**-**B**) Phylogenetic shifts in the scaling relationship between the volume of the ventral lobula (vLO) and the rest of the central brain (rCBR) across 41 Heliconiini species (posterior probability > 0.5). vLO size reductions are identified in *Eueides* and *Philaethria dido*. (**C**) branch specific rates in vLO size plotted against rCBR size.

#### 4.7 Reconstructing pollen feeding evolution

We used three different methods to estimate the discrete trait of pollen feeding at internal nodes in the Heliconiini tree. Both MCMC stochastic character mapping (Figure S10A) and maximum likelihood (Figure S10B) estimates indicate pollen feeding almost certainly arose only once in the Heliconiini, at the base of *Heliconius*, and was secondarily lost in the ‘Neruda’ clade, including *H. aoede*. Maximum parsimony, however, suggests it is equally likely that pollen feeding arose once and was lost once or arose twice within *Heliconius* (Figure S10C).

**Figure S10.**
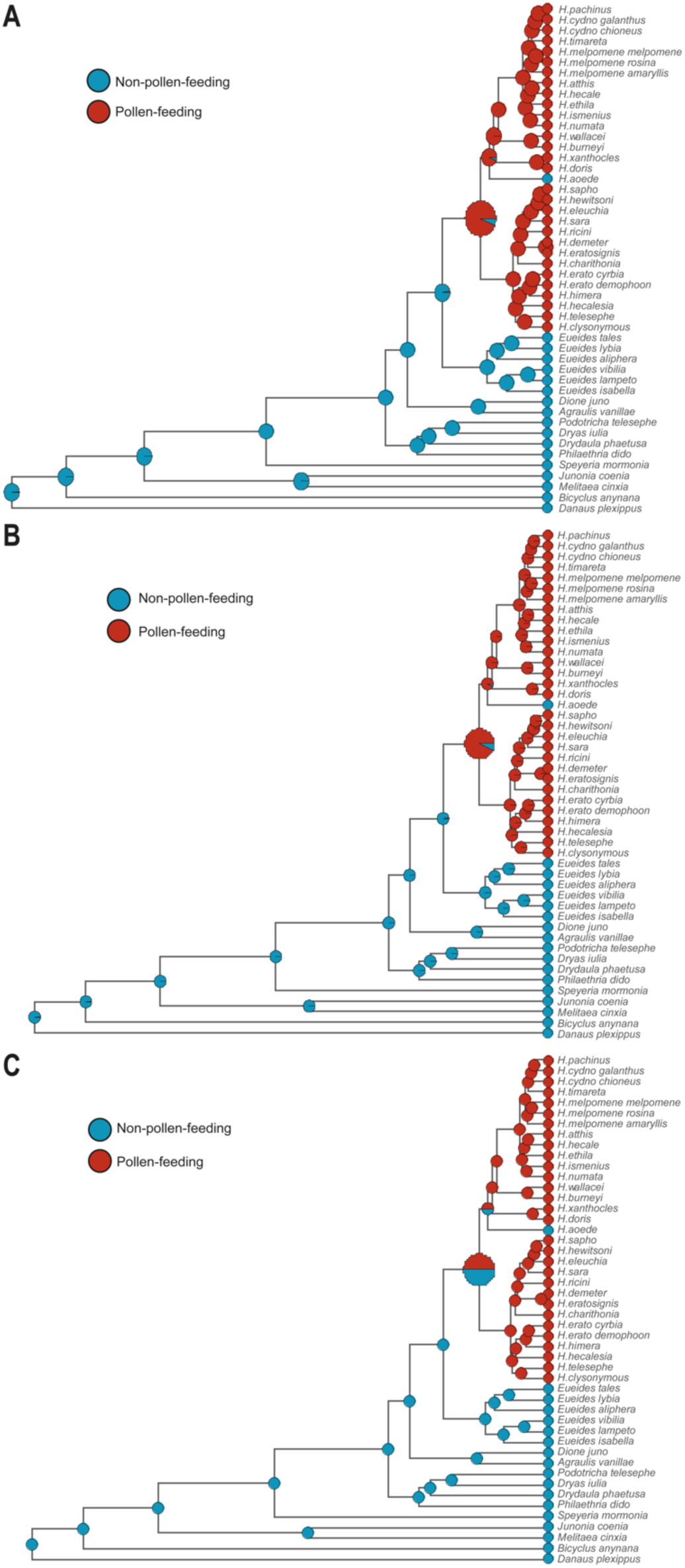
Pollen feeding likely arose once in the Heliconiini, at the base of *Heliconius*. Ancestral state of pollen feeding in the internal nodes of the Heliconiini tree estimated using (**A**) MCMC stochastic character mapping, (**B**) maximum likelihood, and (**C**) maximum parsimony. The colour of the circles represents the probability of pollen feeding at a given node. *Heliconius* root node is enlarged for clarity.

#### 4.8 Visual positive patterning learning in *Heliconius melpomene*

In addition to *Dryas iulia* and *H. erato*, we also assessed the visual positive patterning learning ability in *H. melpomene*, following identical protocols*. H. melpomene* was capable of solving this task (Table S14), with an accuracy comparable to *H. erato* and similarly superior to *Dryas iulia* (Table S14; Figure S11).

**Figure S11:**
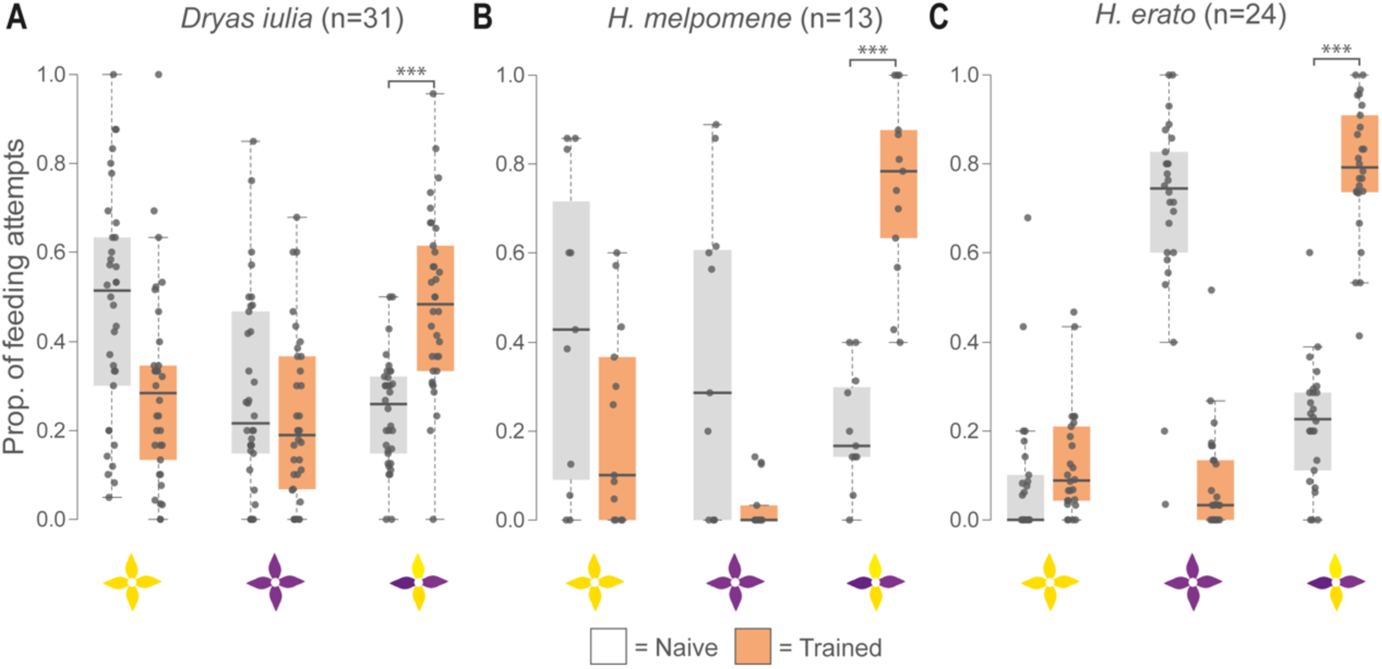
*Heliconius erato* and *Heliconius melpomene* outperform *Dryas iulia* in a visual positive patterning task after eight days’ training. (**A**) *Dryas iulia* was able to solve this visual positive patterning learning task, but its accuracy was lower than both (**B**) *H. melpomene* and (**C**) *H. erato*.

**Table S14.** Visual positive patterning learning in Heliconiini butterflies. Selected pairwise comparisons, corrected multiple comparisons using the Tukey test, between the proportion of correct feeding attempts by *Dryas iulia* (n = 31), *H. erato* (n = 24) and *H. melpomene* (n = 13) before and after training.

| Contrast | Estimate | SE | Z ratio | P value |
| --- | --- | --- | --- | --- |
| *Dryas iulia* naive – *Dryas iulia* trained | -1.018 | 0.111 | -9.195 | <0.0001*** |
| *H. erato* naive – *H. erato* trained | -2.567 | 0.157 | -16.405 | <0.0001*** |
| *H. melpomene* naive – *H. melpomene* trained | -2.593 | 0.270 | -9.610 | <0.0001*** |
| *Dryas iulia* naive – *H. erato* naive | 0.230 | 0.117 | 1.300 | 0.785 |
| *Dryas iulia* naive – *H. melpomene* naive | 0.611 | 0.268 | 2.277 | 0.203 |
| *H. erato* naive – *H. melpomene* naive | 0.381 | 0.285 | 1.336 | 0.765 |
| *Dryas iulia* trained – *H. erato* trained | -1.320 | 0.167 | -7.902 | <0.0001*** |
| *Dryas iulia* trained – *H. melpomene* trained | -0.964 | 0.222 | -4.333 | 0.0002*** |
| *H. erato* trained – *H. melpomene* trained | 0.356 | 0.236 | 1.509 | 0.659 |

